# Expression Patterns of *Mal* genes and its Association with Differential Maltose and Maltotriose Transport rate of Two *Saccharomyces pastorianus* Yeasts

**DOI:** 10.1101/2023.12.06.570446

**Authors:** César I. Hernández-Vásquez, Jorge H. García-García, Esmeralda R. Pérez-Ortega, Adriana G. Martínez-Segundo, Luis C. Damas-Buenrostro, Benito Pereyra-Alférez

**Author notes:** Address correspondence to Benito Pereyra-Alférez.

## Abstract

Microorganisms play a significant role in fermented food biotechnology by converting raw materials in human edible organoleptic and nutritive components, especially in the beer brewing industry. The lager-style beer is the dominant industrial beer type, and it is fermented by *Saccharomyces pastorianus* (Sp) whose members encompass two groups. Typically, strains belonging to group I are deficient in maltotriose consumption. The main variables linked to this phenotype are fermentation conditions, the presence of maltotriose transporters, copy number variation of maltose and maltotriose transporters, and differential genetic regulation. This study was aimed to determine that the differences the alpha-glycoside consumption phenotypes of two Sp strains, Sp820 and Sp790, are related with different phylogenetic distribution and gene expression of the transporters Sc*Mal*x1, Se*Mal*x1, Sc*AGT*1, Se*AGT*1, *MTT*1 and *MPH*x. Biochemical analyses of the transport rate confirmed that the Sp790 strain transported more maltose and maltotriose, 28% and 32% respectively, than Sp820 strain. In addition, detection of Sp790 transcripts indicated the presence of all the *Mal* genes analyzed since the first day of fermentation, whereas Sp820 only presented transcripts for the Sc*Mal*x1, Sc*AGT*1, and *MPH*x genes. These results indicate that a multifactorial phenomenon related with phylogenetic distribution, polymorphisms in transmembrane domains and the difference in the genetic expression of maltose and maltotriose transporters are involved in the phenotypic diversity related with maltose and maltotriose consumption in two lager yeast.

**IMPORTANCE:** Beer is the third most popular beverage around the world and has roughly 90% market share in the alcoholic beverage industry. *Saccharomyces pastorianus* (Sp) strains, which are widely used for lager beer production, have a phenotypic diversity involved in maltotriosa consumption. The fermentation of this sugar is fundamental for the flavor landscape produced during lager beer brewing. This phenotypic diversity encompasses lager yeast strain with remarkable ability to consume maltotriose; Sp group II, to poor capacity of consumption for some lager yeast belonging to Sp group I. Research in this field indicate that variables like conditions of fermentation, presence of maltotriose transporter specific genes, and differential gene regulation can cause this diversity. The significance of our study is to approximate and also contribute to the elucidation of mechanistic variables involved in such phenotypic variability that will allow the development of more controlled and efficient biotechnological processes around beer brewing industry.

## INTRODUCTION

The important role of microorganisms in fermented food biotechnology depends on the enzymatic conversion of the chemical composition of raw substrates to a human edible nutritive product (1, 2). Among the most popular fermented food are beer which is the third most widely consumed beverage around the world and has a considerable market share in the alcoholic beverage industry (3).

In 2018, the global production of beer reached a 191 million kiloliters, up to 0.6% for an increase in comparison with the previous year (https://www.kirinholdings.com/en/newsroom/release/2019/1003_01.html). Since, yeasts are one of the most important components of the brewing process, breweries are dedicated to study the overall characteristics of them in order to improve their process. Typically, the brewing industry uses two types of yeast: Saccharomyces *cerevisiae* (Sc) to produce ale-style beer, whose fermentation temperature ranges from 15°C to 26°C; and *Saccharomyces pastorianus* to produce the lager style beer with fermentation temperatures between 5°C-16°C (4, 5). *S. pastorianus* (Sp), an interspecies hybrid of *Saccharomyces cerevisiae* (Sc) and a cryotolerant yeast *Saccharomyces eubayanus* (Se), is responsible for the world’s best-selling beer, the lager-style (6–8). Nowadays, two distinct types of Sp are recognized based on the chromosome content and sugar consumption. The Saaz-type (group I) and the Frohberg-type (group II), where the main physiological difference is the ability to consume maltose and maltotriose (9). There are two hypotheses that explain the lager yeast’s origin: i) the existence of two independent hybridization events involving different yeasts, Sc and Se (6, 10, 11); and ii) both groups of lager yeast share a single hybridization event, and the evolution resulting from that hybridization shaped genomic divergences between both groups (12). The dominance of Sp in brewing industry suggests that this hybrid has a selective advantage over the parent strains. The key characteristics include faster growth rates in brewing wort, improved biological fitness under stress conditions, and efficient maltotriose consumption (10, 13, 14). The transport of maltose and maltotriose is the limiting factor for effective fermentation (15) since they account for 50-60% and 15-20% of the fermentable sugars in a standard brewing wort, respectively (16, 17). Despite the abundance of maltose and maltotriose in brewing wort, their transport and metabolism are regulated by a carbon catabolite repression mechanism, imposing a sugar consumption hierarchy in which maltose and maltotriose are consumed after glucose is exhausted from the wort (18–20). In this sense, maltose is more easily consumed by most brewing yeasts and hence maltotriose is most abundant in later stages of fermentation when glucose and maltose have been consumed (11, 15, 21, 22).

Different factors have been proposed to approximate the deficient maltotriose consumption phenotypes: i) the stressful conditions of the last days of fermentation, rather than the genetic and biochemical disability of yeast (23), ii) presence or absence of specific transporters for maltotriose such as *AGT*1 and *MTT*1/*MTY*1 (24, 25) and iii) regulatory differences in *MAL* genes (26, 27). In addition, formation of chimeric genes that result in highly efficient maltotriose transporters because of genomic recombination appears as one of the mechanisms that generated variation in these highly related yeast groups (28, 29).

In this study we test the hypothesis about the main factor affecting an efficient fermentation in two lager yeast is related with differential expression pattern of maltose and maltotriose transporters. For this, we focus on a comparative analysis of two strains of Sp, using biochemical approaches where the rate of cellular transport of alpha-glucosides under the same culture conditions was determined. Additionally, we performed bioinformatic analysis of genomic and transcriptomic data to characterize the structure and expression levels of the *MAL loci* (30, 31). Finally, we experimentally validated the bioinformatic analyses by reverse transcription and PCR.

## RESULTS

### Abundance of permeases in the strains studied

The results show the same number of *loci* for *AGT*1, *MPH*x and *MTT*1 permeases in each strain. However, for the *MAL*31 genes, four *loci* were present on Sp790 strain and at six *loci* on Sp820 strain (Table 1 and Fig. S1). Interestingly, both strains had *MTT*1 transporter in their genome, however, the length of their nucleotide sequence was incomplete. The nucleotide and amino acid length of permeases found in each strain varies considerably (Table 1), with only *AGT*1, and *MAL*31 being complete in each genome. Also, the structural distribution of *MAL loci* was different from the canonical MAL *loci*, this canonical structure was observed only at scaffold15 in Sp790 (Fig. S1). This is expected due to the aneuploidy genome features and the sub telomeric locations of these genes may affect the assembly and annotations procedures (11, 32). To validate the completeness of these permeases we used transcriptomic information to obtain the functional product of each transporter (Table 1). The main observation is that both strains have the MAL31p transporter complete in each transcriptome and apparently the MTT1p transporter was not present.

**Table 1.**
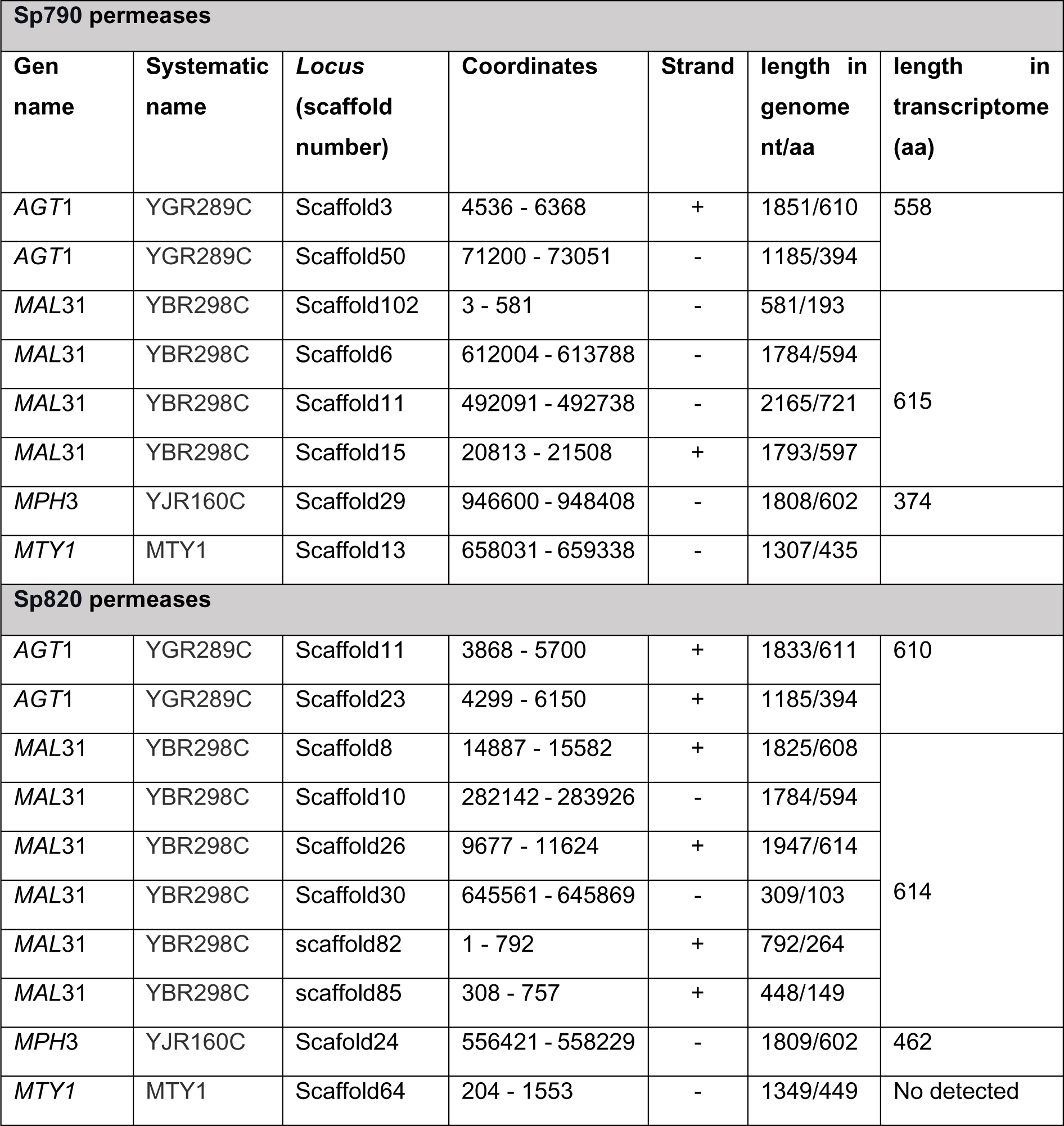
The location and size of permeases for maltose and maltotriose in the studied strains.

| Sp790 permeases |  |  |  |  |  |  |
| --- | --- | --- | --- | --- | --- | --- |
| Gen name | Systematic name | Locus (scaffold number) | Coordinates | Strand | length in genome nt/aa | length in transcriptome (aa) |
| *AGT1* | YGR289C | Scaffold3 | 4536 - 6368 | + | 1851/610 | 558 |
| *AGT1* | YGR289C | Scaffold50 | 71200 - 73051 | - | 1185/394 |  |
| *MAL31* | YBR298C | Scaffold102 | 3 - 581 | - | 581/193 | 615 |
| *MAL31* | YBR298C | Scaffold6 | 612004 - 613788 | - | 1784/594 |  |
| *MAL31* | YBR298C | Scaffold11 | 492091 - 492738 | - | 2165/721 |  |
| *MAL31* | YBR298C | Scaffold15 | 20813 - 21508 | + | 1793/597 |  |
| *MPH3* | YJR160C | Scaffold29 | 946600 - 948408 | - | 1808/602 | 374 |
| *MTY1* | MTY1 | Scaffold13 | 658031 - 659338 | - | 1307/435 |  |
| Sp820 permeases |  |  |  |  |  |  |
| *AGT1* | YGR289C | Scaffold11 | 3868 - 5700 | + | 1833/611 | 610 |
| *AGT1* | YGR289C | Scaffold23 | 4299 - 6150 | + | 1185/394 |  |
| *MAL31* | YBR298C | Scaffold8 | 14887 - 15582 | + | 1825/608 | 614 |
| *MAL31* | YBR298C | Scaffold10 | 282142 - 283926 | - | 1784/594 |  |
| *MAL31* | YBR298C | Scaffold26 | 9677 - 11624 | + | 1947/614 |  |
| *MAL31* | YBR298C | Scaffold30 | 645561 - 645869 | - | 309/103 |  |
| *MAL31* | YBR298C | scaffold82 | 1 - 792 | + | 792/264 |  |
| *MAL31* | YBR298C | scaffold85 | 308 - 757 | + | 448/149 |  |
| *MPH3* | YJR160C | Scaffold24 | 556421 - 558229 | - | 1809/602 | 462 |
| *MTY1* | MTY1 | Scaffold64 | 204 - 1553 | - | 1349/449 | No detected |

To get insight on the variation of the transporters between strains we performed multiple sequence alignments to get pairwise identity matrix, we only included the permeases present in the transcriptome of each yeast strain. Due to annotations and strain specific features not all *Mal* genes alleles and variants were found. We observed transcripts redundancy and whose which share more than 97% of identity was labeled as the same transcript (Fig. S2).

The strain Sp820 has two functional *AGT*1 transporters, TRINITY_DN738_g1_i3 and TRINITY_DN1830_c0_g1_i6 (Fig. 1). The Sp790 has only TRINITY_DN615_c0_g1_i1 transcript annotated as *AGT*1 and share 90% identity with TRINITY_DN738_g1_i3 from Sp820 and 100% identity with TRINITY_DN1830_c0_g1_i6 isoform also from Sp820, indicating that this strain has another functional *AGT*1 variant. We found that *MAL*31 and *MAL*61 transcripts showed sequence differences, especially TRINITY_DN227_c0_g2_i15, TRINITY_DN227_c0_g2_i6 from Sp790 and TRINITY_DN389_c0_g2_i5 and TRINITY_DN389_c0_g2_i2 from Sp820. These trinity isoforms share above 98% identity between them but less than 15% of identity with other MAL31 transporters (Fig 1). Another observation is that TRINITY_DN227_c0_g2_i13 transcript from Sp790 share less than 97% of identity from other MAL31 genes in the same strain and in the Sp820 strain. Also, isoform TRINITY_DN389_c0_g2_i2 annotated as MAL61_2 share 100% identity with isoform TRINITY_DN227_c0_g2_i6 annotated as MAL31 from Sp790 and 98% identity and below with remaining isoforms (Fig. 1). The isoforms annotated as *MPH*x from Sp820 show a very dissimilar distribution of percentage of identity, below 15% with remaining permeases, also, *MPH*x isoforms from both strains have deletions affecting important transmembrane domains (Fig. 1 and Fig. S3).

**Fig. 1.**
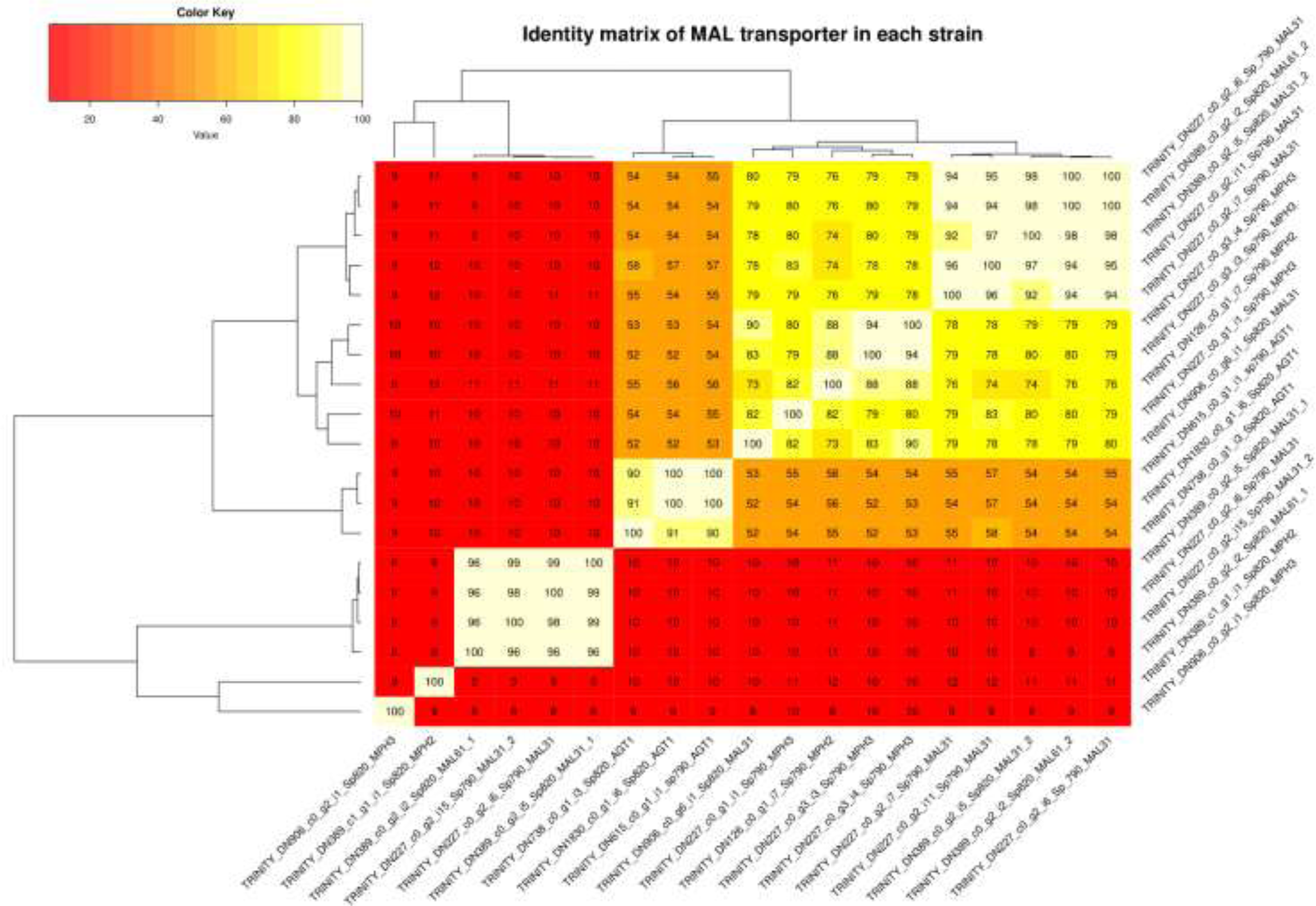
Identity matrix as heat map of paired comparisons of all permeases found in transcriptome data of each strain.

These discordant automatic annotation results prompted us to search for the closest permease reported in the database for the *MAL*31 and *MAL*61 transporters. In Sp820 strain, the isoform TRINITY_DN389_c0_g2_i2, (redundant isoforms in Fig. S2) and the TRINITY_DN389_c0_g2_i5 isoform showed a 100% and 98% identity with a maltotriose transporter (sequence ID: AAY67701.1) (25), representing an interesting observation since this strain has a genomic composition same as group I lager yeast (9). The remaining TRINITY_DN906_c0_g6_i1 showed 99.83% of identity with a maltose transporter reported earlier (ID: QID83234.1) (12).

The strain Sp790 showed the exact behavior of its genes TRINITY_DN227_c0_g2_i15 and TRINITY_DN227_c0_g2_i13 (redundant isoforms in Fig. S2), showing 99.67% and 98.37% identity with maltotriose transporter (ID: AAY67701.1 and CBZ39529.1 respectively). Both strains have over 98% identity with the maltotriose transporter (25) and considering the E-value equals to 0 and 100% of coverage in blastp report, this result indicates that the gene functions as an efficient maltotriose transporter. However, the strain Sp820 displayed a poor maltotriose consumption phenotype relative to Sp790 strain. Because of these conflicting results, we performed multiple alignment analyses to determine the phylogenetic distribution of these transporters under a maximum likelihood algorithm.

### Structural analysis, expression levels and phylogenetic distribution of the permeases

To obtain information about the variation in amino acid sequence and the structure of the *MAL*31 transporter proteins of the studied strains, we made predictions of their primary, secondary, and tertiary structures. First, we performed multiple alignments and predictions of transmembrane domains between the proteins in our database and our *Mal* proteins to determine the sequence and consensus amino acids in the reported transmembrane domains (33).

The *AGT*1 and *MAL*31 transporters of both strains have twelve transmembrane domains (TMD1-12), which is characteristic of MSF transporters (34–36). The *AGT*1_i1 transporter of the Sp790 strain, despite its incompleteness (558 aa), has the twelve canonical transmembrane domains for the MSF family members. The missing amino acids in this protein are related to a region in the intracellular domain that may be associated with the negative regulation of the protein (20, 37). Some transcripts, related with *MPH*x and *MAL*31 genes in both strains, do not show transmembrane domain predictions and some others showed deletions affecting specific TMD structure (Fig. S3). These results were confirmed when 3D model predictions show that TRINITY_DN227_c0_g2_i15 and TRINITY_DN227_c0_g2_i6 from Sp790 and TRINITY_DN389_c0_g2_i5, TRINITY_DN389_c0_g2_i2 from Sp820, those that do not shown TMD predictions, match with MAL regulatory protein 3D structure (Fig. S4). For the remaining non-redundant permeases, the 3D structure prediction matches with alpha-glycoside transporters present in the database, showing a high confidence value between TMD structure and predicted aligned error distributions similar between them (Fig. S4 and Fig. S3). This result allows us to reassign the transcripts that were primary annotated as MAL31 transporters to MALx3 regulatory proteins.

Regarding to the amino acid conservation pattern, we observe the same pattern reported before (33) that show a conservation of E120, D123, E167 and R505 amino acids in 100% of the transporters analyzed here. However, T505 and S557 amino acids are also involved in maltose translocation (38), and these amino acids are not conserved throughout the sequences included in this study. Instead of T505 we observed an Asparagine (N) conserved above 50% in all transmembrane domain 11 (TMD11) sequences (Fig. S5). In the cases of TRINITY_DN615_c1_g1_i1, from Sp790 and TRINITY_DN1830_c0_g1_i5 and TRINITY_DN738_c0_g1_i3 form Sp820 we observed an Isoleucine (I) instead of N505 and T505 (Fig. S5). Lastly, we observed an Alanine (A) conserved above 50% in all TMD12 sequences. However, for the isoforms TRINITY_DN615_c1_g1_i1, from Sp790 and TRINITY_DN1830_c0_g1_i5 and TRINITY_DN738_c0_g1_i3 form Sp820, we observed a Threonine T557 substitution instead of S557 in TMD12 (Fig. S5).

The presence/absence distributions of the maltose and maltotriose transport systems in each strain indicates that strain Sp820 have a larger number of elements because it has another allele linked to *AGT*1 and one more linked to a *MAL*31 gene. Based on the mentioned above, we evaluate the expression levels and the phylogenetic distribution of these transporters in a maximum likelihood context to obtain more information about the phenotypic and sequence differences in maltose and maltotriose transport of these related yeasts.

The general picture of the expression levels of the maltose and maltotriose transporters, show that the Sp790 strain exhibits higher transcriptional activity for all but one transporter, the *MAL*61 showed more expression in Sp820 strain. Specifically, TRINITY_DN615_c0_g1_i1 annotated as *AGT*1 from Sp790 strain show a median FPKM value of 2.6 and 6.08 in the second and fifth day of fermentation, respectively, compared with FPKM = 0.22 and 1.39 from the isoform TRINITY_DN1830_c0_g1_i6 from Sp820 in the same fermentation days (Fig. 3A). Low FPKM values for *AGT*1 are unexpected, as this gene is important for the maltose and maltotriose utilization in ale and lager yeast (39). For *MAL*31 genes we observe that the isoform TRINITY_DN227_c0_g2_i15 from Sp790 strain show the highest expression value in the second and fifth day of fermentation, 329.26 and 180.08 respectively, compared with the highest expressed TRINITY_DN389_c0_g2_i5 isoform in Sp820 strain showing a 34.52 FPKM on the second day of fermentations and TRINITY_DN389_c0_g2_i2 isoform showing 50.12 FPKM on the fifth day of fermentation. As mentioned above, Sp790 strains has two isoforms, TRINITY_DN277_c0_g2_i15 and TRINITY_DN277_c0_g2_i13 that share more than 98% identity whit maltotriosa transporter (as described in above section), and these isoforms are highly expressed compared with Sp820 TRINITY_DN389_c0_g2_i2 and TRINITY_DN389_c0_g2_i5 isoforms that also match with maltotriose transporter (Fig. 3B).

**Fig. 2.**
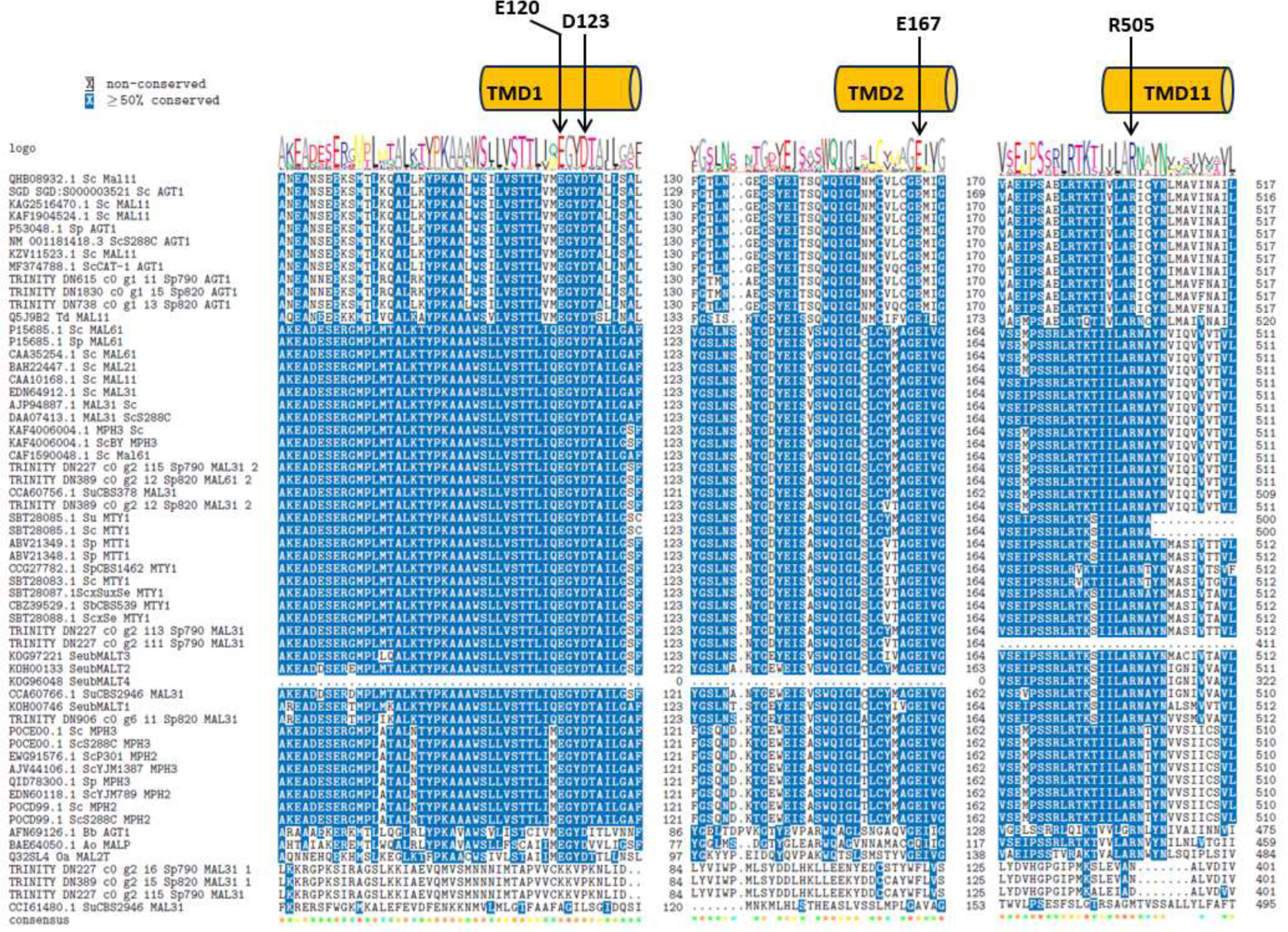
Multiple alignments of maltose and maltotriose transporters. Only the transmembrane domains presenting the important amino acids in the previously proposed translocation mechanism (30) are represented.

**Fig. 3.**
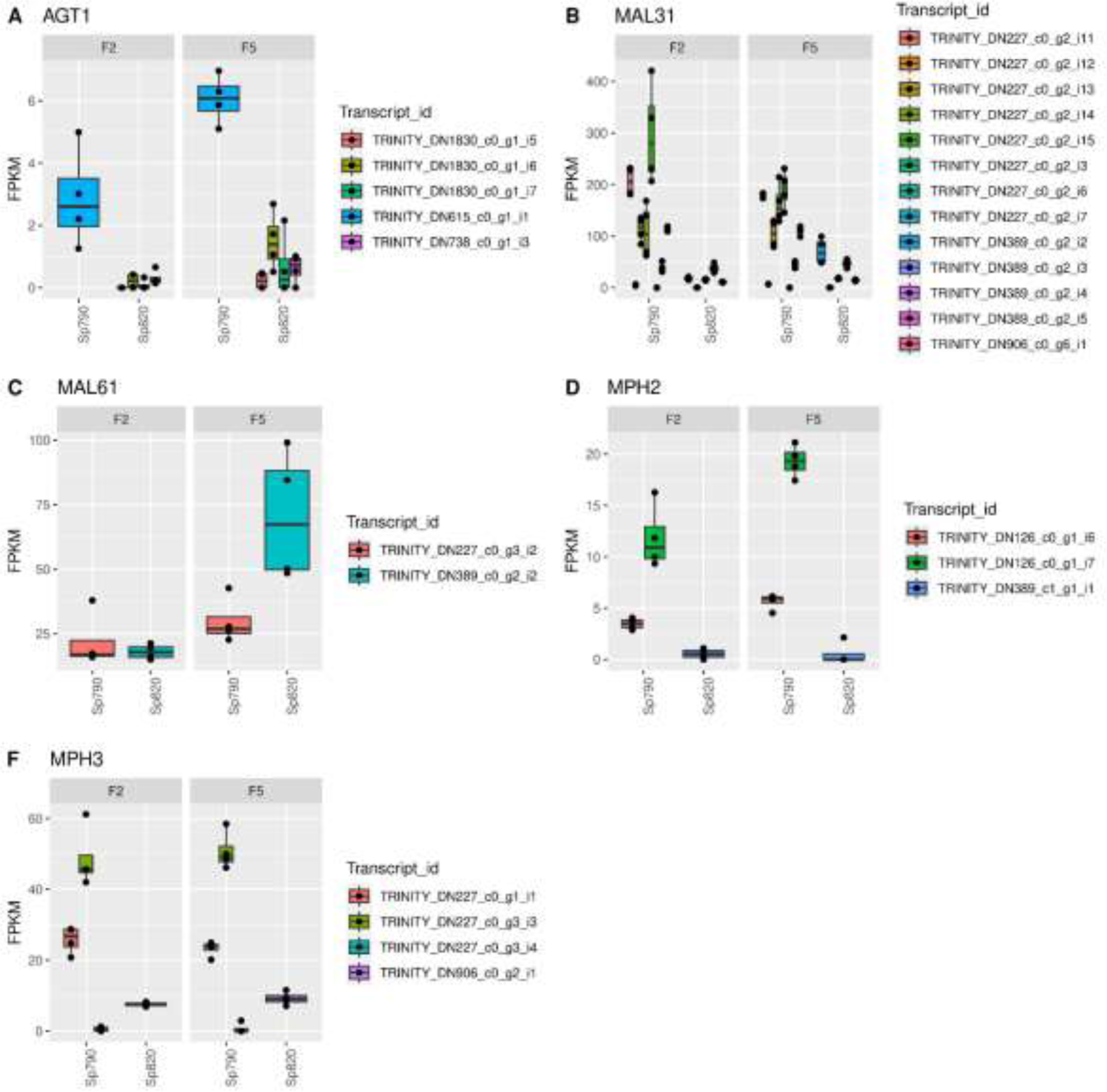
MAL gene expression FPKM values extracted from transcriptomic data of each strain. Note that in **A)** *AGT*1, **B)** *MAL*31, **D)** *MPH*2 and **F)** *MPH*3 genes the strain Sp790 has more expression than the Sp820 strain. The strain Sp820 has higher expression of **F)** *MAL*61 transporter only in the fifth day of fermentation. Transcript id represents all specific transcript assembled for each strain.

Regarding the *MAL*61 transporters, we observed that the transcript isoform TRINITY_DN389_c0_g2_i2 from Sp820 has the higher expression, 67.27 FPKM, on the fifth day of fermentation. In the second day of fermentation, we observed the same transcriptional oscillation between the strains (Fig. 3C). Finally, the transcriptional activity of *MPH*2 and *MPH*3 show higher values in Sp790 than in Sp820 (Fig. 3D and 3F), however, all transcript isoform from both strains showed a deletion that affect a considerable number of transmembrane domains affecting the functionality of this transporters (Fig. S3).

Additional to the different expression behavior of MAL transporters that show more expression activity in Sp790 than in Sp820, the phylogenetic distribution also differs between these strains. In general, we observed that the TRINITY_DN227_c0_g2_i7, annotated as *MAL*31 transporter, has redundancy with TRINITY_DN227_c0_g2_i3/i13 (Fig. S2), from Sp790 has a similar phylogenetic behavior as in MTT1/MTY1 transporter from the database. In contrast with all transcript isoforms annotated as *MAL*31 genes, observed in Sp820 (Fig. 4). This result strongly suggest that the Sp790 strain has a highly efficient maltotriose transport system due to the *MTT*1 transporter activity (25, 40). And in contrast, the Sp820 strain lacks an *MTT*1 gene and the pairwise comparisons results described above were an artifact output. However, this scenario shows conflicting information and requires physical confirmation.

**Fig. 4.**
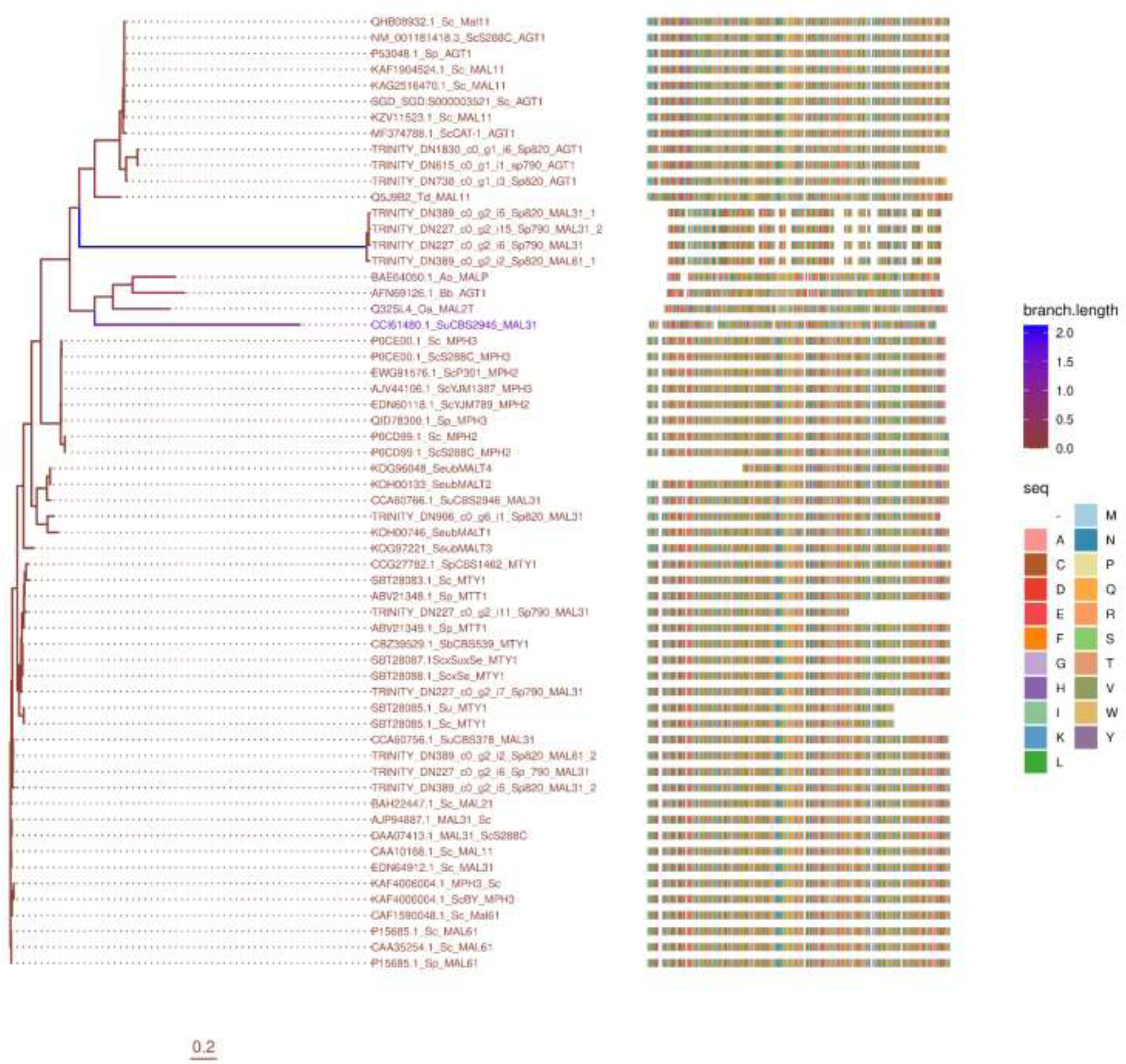
Maximum likelihood phylogeny for the studied permeases and those reported in the databases. Note how the isoforms i11 and i7 of annotated MAL31 permease of the Sp790 strain groups with the *MTT*1/*MTY*1 permeases reported in Sp. It is also important to note that the permeases of the yeast Sp820 has a different phylogenetic distribution from *MTT*1/*MTY*1 genes.

### Global expression analysis

We performed differential gene expression analysis using the *Saccharomyces pastorianus* CBS 1483 genome as a reference. The *MAL* loci analyzed in this work showed 99-100% identity and a coverage of 87- 100%, with chromosomal regions of the mentioned strain (Table S1). In the context of the general transcriptomic response, we only focused on maltose and maltotriose transport differentially expressed genes. In the second day of fermentation comparisons between both strains (contrast groups F2_820 vs F2_790), we observed two differentially expressed genes in strain Sp820; *MAL*11_2 and *MAL*11_1 (Fig. 5) which in *S. pastorianus* represent a specific allele called *AGT*1 with considerable affinity for maltotriose Km 18.1 mM (41). According to the analysis of the regulatory region of the *AGT1* gene (Table S2), the allele encoding this protein has six binding sites for the transcription factor *MAL*63, an activator of the *MAL* genes, two of them at a distance of - 2600 and one at −1000 bp from the transcription start site. Using the previously established model, insertions in the regulatory region separated the *MAL*63 sites to the −700 to −800 bp position, resulting in reduced expression of the *MTT*1 gene (26). We can establish functional relationships between *AGT1 (MAL*11) alleles expressed at the second day of fermentation (F2) for Sp820, because the functional allele encoding the complete 610 aa protein has two *MAL*63 binding sites at −375 and −95 bp position, the positional range required for efficient transcription.

**Fig. 5.**
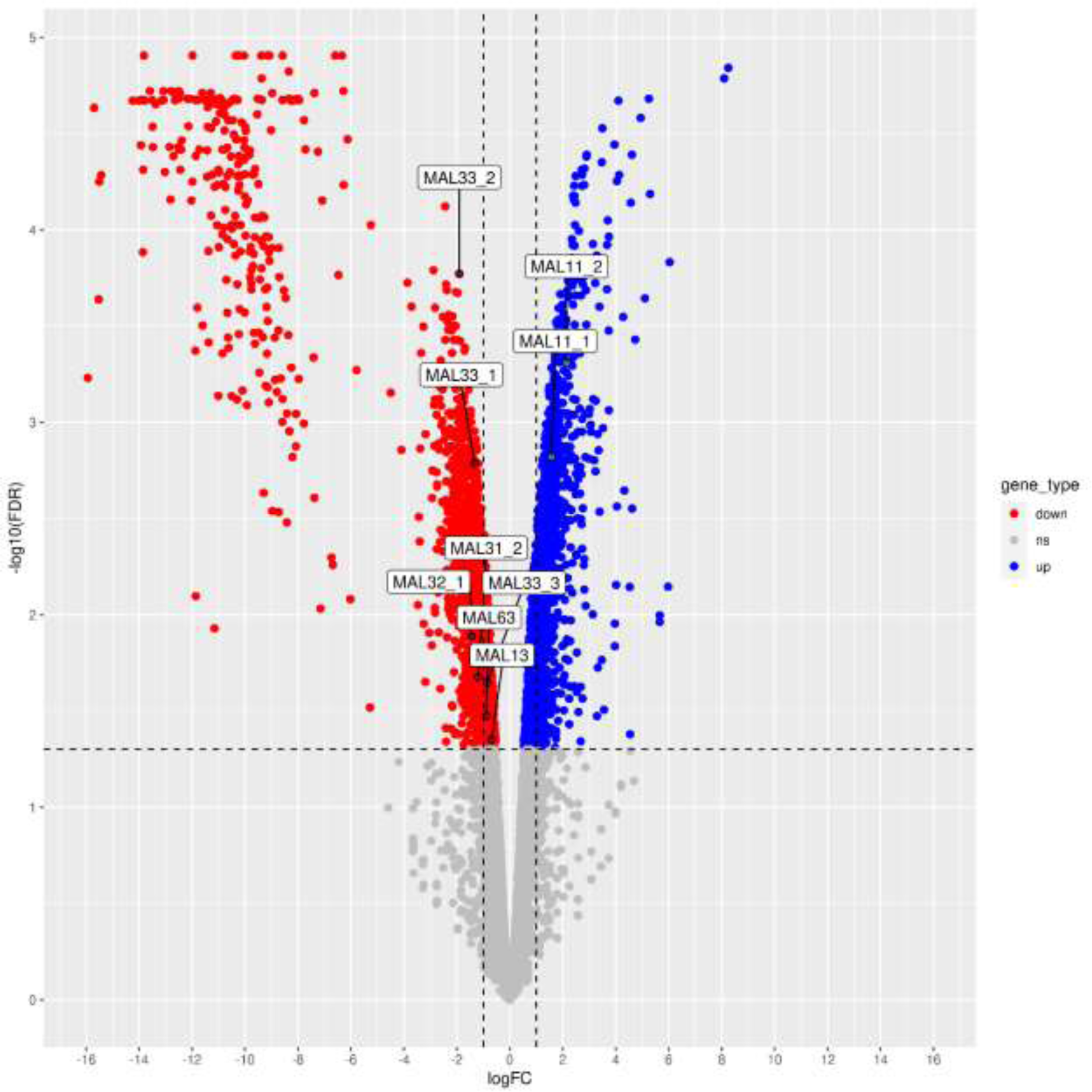
Volcano plot showing genes differentially expressed genes in the second day of fermentation, genes in blue show higher expression in strain Sp820, genes in red lower expression, and genes in grey are not differentially expressed. Red and blues genes are inversely interpreted for Sp790.

On the other hand, the *MAL*33_1, *MAL*33_2, *MAL*32_1, *MAL*31_2, *MAL*33_3, *MAL*63, and *MAL*13 genes are differentially expressed in strain Sp790 (Fig. 5 and Table S4). These genes may also perform regulatory functions: *MAL*33_1-3, *MAL*63, and *MAL*13. The high number of these genes may be related to the efficiency of strain Sp790 in consuming maltose and maltotriose. This is consistent with the regulatory region analysis, as the strain Sp790 has almost the same number of *MAL*31 genes and has similar pattern in *MAL*63 binding sites as observed in Sp820 (Table S3). Also, the transcriptional activity, measured as FPKM, showed that Sp790 has considerably more expression for the transcript isoforms annotated as *MAL*13 and *MA*L63 than the observed FPKM values for Sp820 (Fig. 3 and Fig. S6). This is expected as is more likely that a regulatory network evolves by means of mutation in cis-trans fashion in yeast (42).

The same pattern of expression is seen on the fifth day of fermentation (F5), where strain Sp820 shows expression of maltose and maltotriose transporters, *MAL*11 (*AGT*1) and *MAL*31, as well as some maltases such as *MAL*32 and *MAL*62 (Fig. 6 and S table 5). This is particularly interesting since the functional allele of *AGT1* has been held responsible for efficient fermentation in early publications (15, 17, 43); however, the affinity of *AGT1* for maltose is greater than for maltotriose (5-17 mM and 18 mM, respectively). Competition events may be taking place in which maltose transport take precedence, as suggested by the observation on the second day of fermentation analytical data (F2), where maltose concentration is more than twice of maltotriose (4.82 ± 0.61% vs 2.16 ± 0.26% w/w) (Fig. 7C). This could explain the poor maltotriose transport efficiency of Sp820 strain. Similarly, strain Sp790 has more regulatory genes *MAL*63, *MAL*33_1-3, and some transporters such as *MAL*11_3 and *MAL*31_4. Considering the higher number of predicted positive regulators (*MAL*x3) and the MAL63 binding sites in the regulatory regions of the *MAL* genes of strain Sp790 (Table S3), and the presence of the *MTT*1 gene (Fig. 2), this represents a combination of two critical variables important for efficient fermentation: differential gene regulation events and presence of maltotriose specific transporters.

**Fig. 6.**
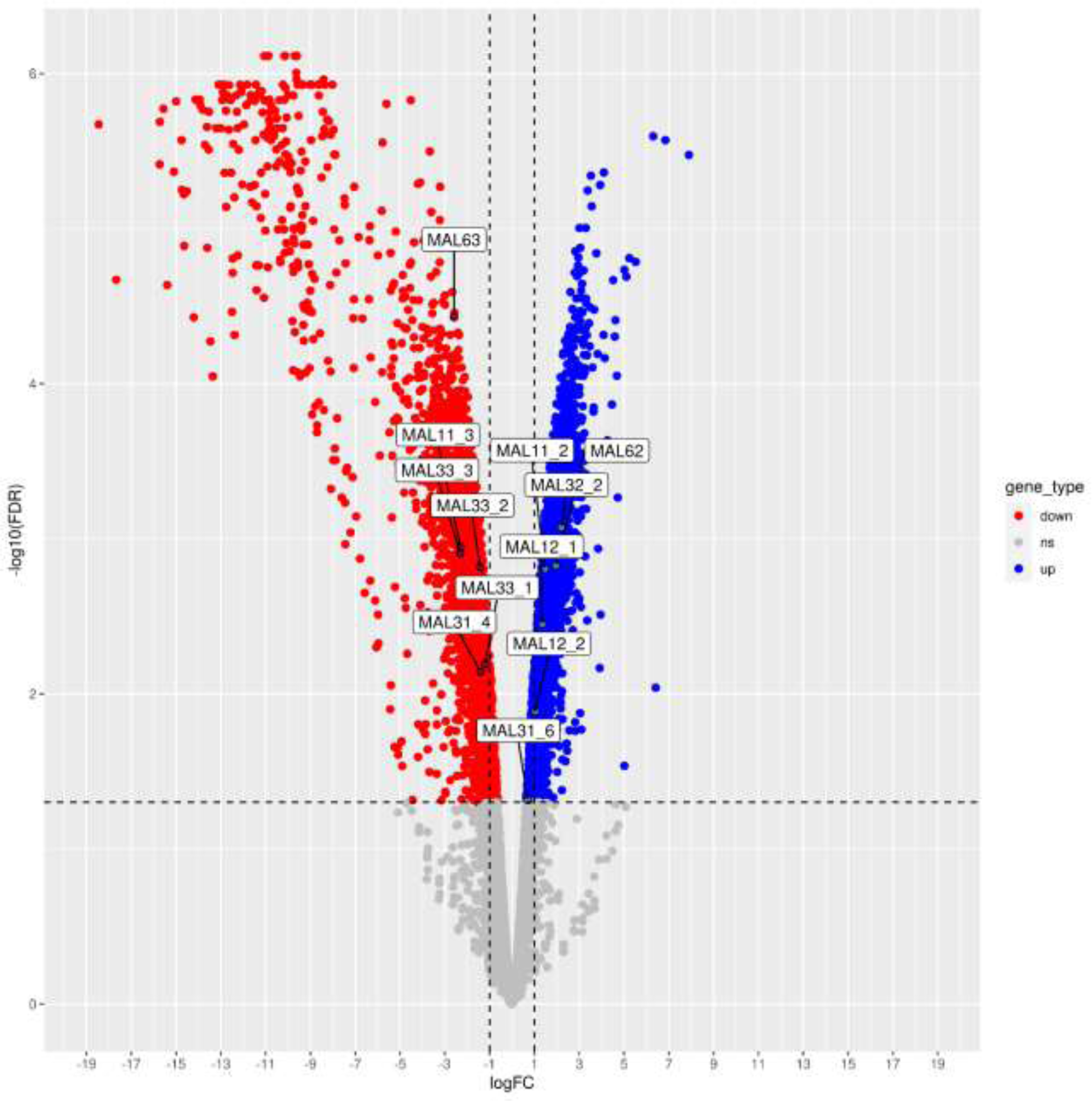
Volcano plot showing genes differentially expressed in the fifth day of fermentation genes in blue show higher expression in strain Sp820, genes in red lower expression and genes in grey are no-differentially expressed. Red and blues genes are inversely interpreted for Sp790.

**Fig. 7.**
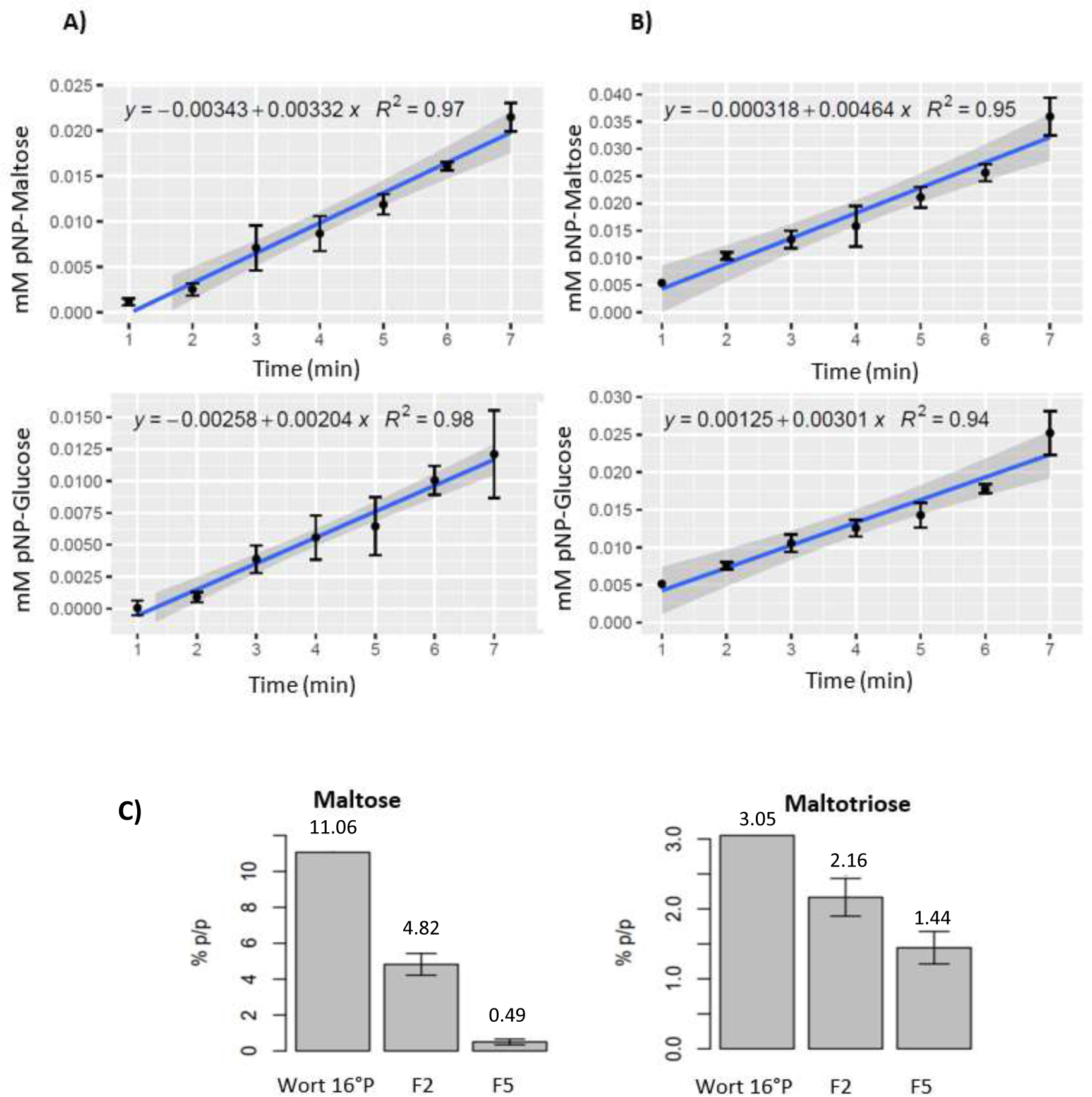
Transport of pNP-α-glucose or pNP-α-Maltose in strains Sp820 and Sp790. **A**) Transport plots for pNP-α-Glucose (alpha-glucoside structurally related to maltose) and pNP-α-maltose (alpha-glucoside structurally related to maltotriose) in strain Sp820. **B**) Transport graphs of the same pNP- α-glucosides in strain Sp790. **C**) Maltose and residual maltotriose on second and fifth day of fermentation for strain Sp820. Data for strain Sp790 was undetectable.

The previously proposed model suggests that a greater number of binding sites at *MAL*63 indicates increased expression of downstream genes (27) and that the distance from the transcription start site of this promoter and regulatory the site is related to expression efficiency (25) and may explain the higher maltose and maltotriose transport efficiency observed in strain Sp790.

In addition, the null detection of fermentable sugars at the second (F2) and fifth (F5) of fermentation day in strain Sp790 may provide a functional example of differences in gene regulation. This is reinforced by the observation that strain Sp790 expresses transcripts for the *MTT1* gene, which encodes a very efficient maltotriose transporter (Km 16 mM) (25, 40). Finally, the presence of the *MPH*3 gene may not have functional implications for the transport of maltose and maltotriose, as the analysis of its regulatory region found only one *MAL*63 binding site for both strains, and reduced FPKM and no statistical evidence was found for *MPH*x genes. This is consistent with the results of the previous report, in which a low or null number of copies of this gene was observed, deprecating the role of *MPH*3 in efficient fermentation (24).

### Biochemical analysis of the transport rate of maltose and maltotriose in the study strains

We determined the pNP-α-glucose and pNP-α-maltose transport curves for the strains Sp790 and Sp820. chemically labeled pNP substrates are transported into the cell by alpha-glycoside transporters (*Mal*x1, *AGT1*, *MPH*x, and *MTT1*), where alpha-amylases hydrolyze the alpha bond of maltose and maltotriose, releasing the pNP. After permeabilization and pNP extraction, spectrophotometric measurements were performed at 500 nm for quantitative determination (44). Since the stoichiometric ratio of pNP to the alpha-glucoside of interest is 1:1 (45), it can extrapolate the concentration of transported pNP to the concentration of labeled alpha glycoside. In general, we observed that maltose transport is more efficient than maltotriose transport in both strains (Fig. 7A and 7B).

it is reported that the hierarchical preference of consumption of carbon source is glucose> maltose> maltotriose (18, 19). In addition, transporters such as *AGT*1 and *MAL*x1 have a higher affinity for maltose (46). This explains that the transport of maltose (pNP-α-Glucose; glycoside structurally related to maltose) is naturally more efficient than the transport of maltotriose (pNP-α-Maltose; glycoside structurally related to maltotriose). When we compared the transport of the same carbohydrate in different strains, we found that the strain Sp790 had a higher consumption rate of the respective carbohydrate. Specifically, the Sp790 strain exhibits a transport rate of 282 nmol min^-1^ mg^-1^ dry weight for maltose. Compared to the result for the Sp820 strain for the same carbohydrate, 202 nmol min^-1^ mg^-1^ dry weight (Fig. S7). These results are exciting because the genomic and transcriptomic analysis showed that both strains have the genes involved in maltose transport (*AGT1*, *MAL*x1, and *MPH*x), suggesting that the genes associated with maltose transport have a differential transcript quantity that may be mediated by differences in genetic regulation or copy number variation, as the Sp820 strain has two functional alleles of the *AGT*1 transporter. In this context, genetic regulation in ale and lager yeast are responsible for the different phenotypes in the consumption of maltose and maltotriose. Specifically, the regulatory region of the *AGT*1 transporter in ale yeast has two insertions that result in an additional binding site for *MAL*x3, a positive regulator of *MAL* genes, and multiple binding sites for *Mig*1, a repressor of *MAL* genes in the presence of glucose (27, 47). These regulatory changes could be involved in the different phenotypes observed for the rate of maltose transport in the strains studied.

The same scenario applies to maltotriose transport. The transport rate of this carbohydrate was higher in Sp790 than in Sp820, 183 nmol min^-1^ mg^-1^ dry weight versus 124 nmol min^-1^ mg^-1^ dry weight, respectively. However, the reports of differences in maltotriose transport suggest the existence of more specific maltotriose transport systems. Two independent studies have identified the presence of a transporter, *MTT*1/*MTY*1, that has a higher affinity for maltotriose, km 16-27 for maltotriose and km 61-88 for maltose (25, 40). This hypothesis of more efficient transport systems for maltotriose is very attractive since, despite the physiological classification of type I and type II lager strains, where type I strains show deficiencies in maltotriose consumption (9), the presence of this transporter is responsible for an efficient fermentation phenotype in a group I strain of *S. pastorianus* that closely resembles the fermentation efficiency of group II yeasts (24). However, the latent doubt that the Sp820 strain may possess the *MTT*1 is contradictory, since the TRINITY_DN227_c0_g2_i2 and TRINITY_DN389_c0_g2_i5 transporter shows an identity of more than 98% with the *MTT*1 transporter in the database and in contrast to the distinct phylogenetic distribution far from *MTT1* clusters. In view of these results, we decided to perform a genomic and qualitative transcript detection by reverse-transcription and PCR to complement these outcomes.

### Molecular detection of permease genes and transcripts in each strain

To experimentally validate our bioinformatic projections, we performed detection of the genes and transcripts of permeases in each yeast by PCR and retro-transcription-PCR. For this purpose, we use previously reported primers to discriminate allelic variants of *AGT1* (Sc*AGT1* and Se*AGT1*), *MAL*x1 (Sc*Mal*x1 and Se*Mal*x1), *MTT*1/*MTY*1 and *MPH*x (24). First, we performed genomic detection of each gene in the study strains. In the process, we noticed two critical results (Fig. 8A): i) both strains generally have *ScMal*x1, *SeMal*x1, *ScAGT1*, *SeAGT1*, *MTT1*, and *MPH*x genes in their genome, and ii) the strain Sp820 has the *MTT*1 gene in its genome, which was confirmed by our bioinformatic pipeline in the genome sequencing data. Finally, we performed detection of the transcripts of each permease in the first two days of fermentation. Using these data, we noticed two important observations (Fig. 8B): i) both strains show transcription of all permeases analyzed. And more interestingly, the Sp820 strain shows the expression of the *MTT*1 gene at the second day of fermentation. And ii) the permeases showed differences in regulation: On the first day of fermentation, the Sp790 strain showed all permeases analyzed in the experiment, in contrast to the Sp820 strain, which showed only the *ScMal*x1, *ScAGT*1 and *MPH*x permeases on the same day.

**Fig. 8.**
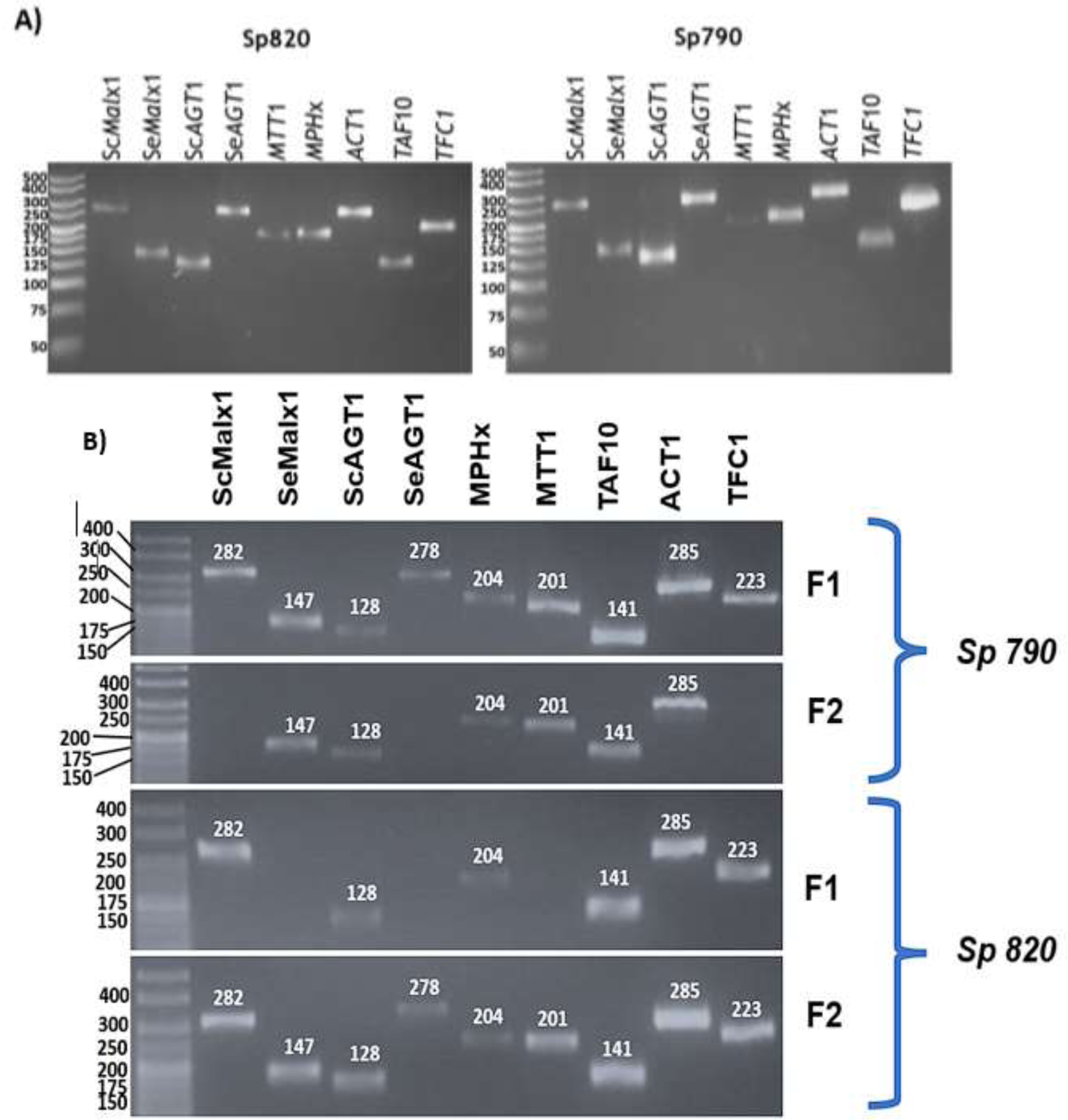
Molecular detection of maltose and maltotriose permeases in each strain analyzed. **A**) Genomic detection of permeases. **B**) Detection of the transcripts on the first and second day of the fermentation of the studied strains. Notice that on the first day the strain Sp790 shows all transcripts for Mal genes and on the same day the strain Sp820 only shows *ScMal*x1, *ScAGT*1 and *MPH*x.

On the second day of fermentation, the Sp790 strain showed only *SeMal*x1, *ScAGT*1, *MPH*x, and *MTT*1 permeases. In contrast, the Sp820 strain showed expression of all permeases analyzed. These data support the bioinformatic inferences regarding the differences in the expression profiles of the analyzed strains and suggest that genetic regulation is the predominant variable in the phenotypic variation of maltose and maltotriose transport in our two analyzed strains.

## DISCUSSION

### The structural variability accounts in the phenotypic variation in alpha-glycoside transport

The observed diversity of proteins is the product of random mutational processes and natural selection, which in turn is responsible for maintaining or enhancing a given phenotype. This overly robust process of protein selection allows 50-80% of residues to be altered without affecting the protein structure (48). However, such changes at specific residues can affect the structure of the protein, which happens more often in regions related to the substrate binding site (49). In this section, we use the previously proposed bioinformatics approach (33) to observe possible specific variations in amino acids closely related to the maltose translocation mechanism. Site-directed mutation experiments have been determined that the critical amino acids in this mechanism are E120, D123, E167, and R504 (33, 45). In this study, we observed a conservation pattern in the above amino acids.

On the other hand, other residues such as T505 and S557 have been associated with the efficiency of maltotriose transport in the *AGT1* gene (38); however, we observe a substitution in these amino acids for all protein sequences analyzed in this work (Fig. S3). For all *AGT*1 we observed an I505 and for all MAL31 transporter N505 rather than T505. For S557 amino acid we observed T557 instead S557 in all *MAL*11 genes. And for *MAL*31 and *MPH*x genes we observed A557 instead of S557. Lastly, it has been determined that a polymorphic discrete sites in TMD 7 and 11 dictate the substrate specificity in *MAL*61, *MTT*1 and *MTT*1- like transporters present in Ale yeast. Specifically, the amino acids T378, N383, A506 and S506 are involved in this specificity (50). Our result indicates that only the isoform TRINITY_DN227_c0_g2_i7 form Sp790 has the T379 and N389 residues that confers maltotriosa specificity (Fig. S8). The remaining transporters and more importantly the transcripts isoforms from Sp820 TRINITY_DN389_c0_g2_i2 and TRINITY_DN389_c0_g2_i5 that matches 98% with maltotriose transporter have the A379 and Y389 substitutions indicating that these transporters are maltose specific (50).

By these observations we can speculate that discrepancies in the above amino acids could affect in maltotriose transport efficiency but underlying high transcription in Sp790 culminates in no detectable residual maltose and maltotriose. But in Sp820 these substitutions may play accumulative factor, combined with differential *MAL* gene transcription relative to Sp790, in its less efficient maltose and maltotriose phenotype.

In general, these results suggest that the industrial conditions of brewing fermentation function as a selective force limiting the population of maltose and maltotriose transport proteins, resulting in less variability in this phenotype. However, this is not an absolute event, as the structural statics of these permeases may be compromised due to their chromosomal location and the hybrid nature of *Saccharomyces pastorianus* (30, 51, 52).

### The variables of expression and identity of the transporters efficiently explain the phenotypic variation in the transport of maltose and maltotriose in the analyzed strains

The *S. pastorianus* strains analyzed in this study showed no fundamental differences in the structure of the maltose and maltotriose transporters analyzed (Fig. 2). However, we observed differences in the expression levels of the permeases and a significant difference in the global expression of the *MAL* genes under the same fermentation conditions (Fig. 3, Fig. 5 and 6, respectively). We speculate that gene regulation networks are the fundamental difference between the two *S. pastorianus* strains. As noted earlier, the high degree of similarity between the proteomes of related organisms supports the idea that changes in gene regulation play the central role in evolution (53). In this case, the genes for transport of maltose and maltotriose in two strains of *S. pastorianus* show variations in their genetic expression. This event is consistent with the current paradigm that a small number of genes and their regulation shape the physiology of higher organisms (54). In this context, it is important to note that the evolutionary origins of lager yeasts are presented as a breeding ground for divergence events in gene expression. Currently, polyploidy (12, 52) and the location of *MAL* genes in the sub telomeric regions of chromosomes (55) are thought to be key factors in adaptation to environments such as beer fermentation.

Recently it was found that cross-linking occurs in the regulation of maltotriose uptake in sub genomes (Sc x Se) in yeast *S. pastorianus* (56). The polyploidy of *S. pastorianus* yeasts enhances the evolutionary process through divergence in gene expression and promotes gene duplications (51). Moreover, research has shown that the difference between Sc and Sp strains is in the regulatory region that controls the expression of some *MAL* genes (26, 27).

In terms of the phylogenetic distribution, the Sp790 strain show that at least one of their transcripts, TRINITY_DN227_c0_g2_i7, clustered with *MTT*1/*MTY*1 transporters evidencing that this strain has an efficient maltotriosa transport system. For the Sp820 strain, no *MTT*1/*MTY*1 was observed and also the polymorphism distributions of TMD7 (50) indicates that the matching of their transcripts, TRINITY_DN227_c0_g2_i7 and TRINITY_DN389_c0_g2_i5, with *MTT*1 was an artifact output.

With this observations, the *MTT*1 gene present in the Sp790 strain supports the previously proposed hypothesis that this gene is responsible for efficient fermentation (21).

As the presence of *MTT*1 in strain Sp820 may be questioned, the variation in the phylogenetic, TMD7 polymorphism distributions and the differential gene regulation in the investigated lager yeast strains, show a clearly multifactorial phenomena involved in the differential maltose and maltotriose uptake observed in these yeasts.

### Under the same conditions, the yeasts show differences in their maltose and maltotriose transport rates

To complement the bioinformatic information obtained so far indicating that both strains possess a gene annotated as *MAL*31, which is more than 98% identical to *MTT1*, we determined the cellular transport rate of pNP-glucose (structural analog of maltose), and pNP-maltose (structural analog of maltotriose) (33, 38, 44, 57). We assumed that we would observe the same transport rate under the same standard reactions and culture conditions because the strains generally have the same maltose and maltotriose transport system. However, we found that the Sp790 strain has an additional 28% and 32% transport for maltose and maltotriose, respectively, compared to the Sp820 strain. These results suggest that if the scenario in which the strains have the same transport system for maltose and maltotriose is true, another variable, such as copy number variation and the different gene regulation, is the underlying phenomenon for the divergence of these phenotypes.

Despite the different industrial conditions used in the strains studied, we found that this lager yeast exhibited fundamental differences in transport rate under the same standard conditions. This suggests that, in contrast to previous studies, fermentation conditions make little or no contribution to the alpha-glucoside transport phenotype (23).

In the context of genetic regulation imparted by catabolite repression and the previously transporter affinity, we can reevaluate the importance of maltose rather than maltotriose transport. Because the *AGT1* gene is responsible for transport of maltose and maltotriose and has different affinities, the affinity for maltose, Km = 14 mM is on average (41, 58–60) greater than that for maltotriose Km = 27 mM. We can predict events of substrate competition in which maltose is taken up before maltotriose. As we can observe from the residual maltose and maltotriose on the second and fifth day of fermentation for the Sp820 strain (Fig. 7C).

The yeast Sp820 transported 56% of the maltose at the second day of fermentation. In comparison, the maltotriose transport at the same day was only 29%. Similarly, at the fifth day of fermentation, yeast Sp820 transported 96% maltose and only 53% maltotriose. This biochemical information may indicate that substrate competition is occurring, and the possible contribution of the additional transcriptional load, observed in Sp790 bioinformatic data, may contributing to the differences in consumption of maltose and maltotriose.

### Physical evidence of permeases suggests differential regulation among yeasts studied

The bioinformatic data showed discrepancies regarding the identity of the *MAL*31 genes of strains Sp790 and Sp820 (Fig. 4). We decided to perform molecular detection of these genes using previously reported primers (23). First, the events observed in the genomic detections (Fig. 8A) indicate that both strains contain the same molecular machinery (Sc*MAL*x1, Se*MAL*x1, Sc*AGT1*, Se*AGT1*, *MTT1*, and *MPH*x) for uptake of maltose and maltotriose from the wort. Although the *MTT*1 gene in the Sp820 strain could not be detected by bioinformatics approaches, the variable related to divergence in gene expression became relevant in these analyzes. However, since genomic screening only indicates the possible phenotypic potential of our yeasts, we decided to screen the transcripts associated with the above genes. In these results, we find that in Sp790 strain all genes are induced on the first day of fermentation despite the presence of glucose in the wort. In contrast, in Sp820 strain only the Sc*MAL*x1, Sc*AGT1* and *MPH*x genes are present. This is consistent with the residual amounts of maltose and maltotriose present in strain Sp820 on the second and fifth days of fermentation (Fig. 6C), as the Sc*AGT1* allele was previously found to be nonfunctional in the Sp strains (40, 61) and the *MPH*x gene was excluded as important for alpha glucoside consumption (24). At the second day of fermentation, strain Sp820 has all the genes analyzed, but probably the observed repression caused a delay in the consumption of the maltose and maltotriose from the wort (Fig. 6C).

Under these circumstances, it is reasonable to assume that strain Sp790 diverged in the genetic regulatory networks related to the hierarchy of sugar consumption in the brewing wort to culminate in one of the most important phenotypes of the industrial niche: efficient consumption of alpha-glucosides. While strain Sp820 still exhibits adaptation events that affect its transport efficiency of alpha-glycosides. If we position these two yeasts at different times in an evolutionary pathway, it is possible that the Sp820 strain will increase its transport efficiency of alpha glycosides and its overall fermentation profile through assisted evolution (62, 63).

### The *Mal loci* pose problems for approximation by next-generation sequencing

Many efforts have been made to mitigate the limitations of the new sequencing technologies in resolving repetitive and conflicting regions in the genomes of some yeasts. Some reports avoid the use of such technology, opting to use hybridization experiments (24). Other approaches use long- and short-read sequencing experiments to achieve the highest possible resolution of some lager yeasts (12). Nonetheless, complementing bioinformatics approaches with the transcriptome of interest allows for even higher resolving power of the omics information in question. Furthermore, physical verification of bioinformatic hypotheses must be presented when conflicts like misannotations or transcript misrepresentation (64, 65) as they did in this study. Our bioinformatics approach using short-read sequencing, both in the genome and in the transcriptome, allowed us to establish that variation in phylogenetic distribution and gene expression is extremely important for the alpha-glycoside uptake phenotype. However, one of its limitations is that it showed inconsistent results and poor resolution with respect to the presence of the *MTT1* gene in the Sp820 strain. Here we see the importance of a biochemical and molecular approach to clarify and corroborate our bioinformatic observations.

### Conclusion: Difference in genetic regulation plays a significant role in the variability of Maltose and Maltotriose transport in both strains of *Saccharomyces pastorianus*

The present work sheds light on the factors underlying different phenotypes of maltose and maltotriose consumption that exist in some brewing yeasts. We found that polymorphic regions observed in TMD7 and TMD11 may be involved in maltotriose specificity. Also, under the same fermentation conditions, the yeasts show differences in the expression levels (FPKM values) of the analyzed *MAL* genes and global differential expression, with strain Sp790 showing greater regulatory activity related to the positive activation of the *MAL* genes. We also observed that strain Sp790 exhibited a higher cellular transport rate of pNP-glucose and pNP-maltose than strain Sp820 under the same standard conditions. In addition, through genomic and transcript detection, we confirmed the bioinformatic observations regarding the differences in genetic expression between the two yeasts. Furthermore, we resolved the conflicting information about the identity of the *MTT*1 genes in Sp820. This is an interesting result because the presence of the *MTT*1 gene isn’t enough for efficient alpha-glycoside uptake; it also requires copy number variation and/or efficient transcription levels. Our observations suggest that the main event associated with the differences in the consumption of alpha-glucosides in the yeasts studied here is the differential genetic expression of the transporters for this substrate. However, we cannot exclude that the factor related to the differences in copy number variation of the transporter genes contributes to this divergence in transcriptional load. For this reason, quantitative analysis related to differences in transporter copy number and transcript levels are necessary. Both yeasts have the same machinery for efficient fermentation, but from an evolutionary perspective, yeast Sp790 has produced an efficient alpha-glucoside consumption phenotype. This suggests that yeast Sp820 could achieve efficient fermentation and change in its overall fermentation landscape through an assisted evolutionary experiment.

## MATERIALS AND METHODS

### Strains

Strains identified as Sp820, and Sp790, members of group I and group II respectively, were kindly provided by Cervecería Cuauhtémoc Moctezuma (Monterrey, NL. México). Yeast strains were inoculated into YP-M medium (1% yeast extract, 2% peptone, and 2% maltose) and after 48 hours or when they reached stationary phase, 2 mL aliquots of YP-M medium containing 40% glycerol was prepared in sterile vials and stored at −20°C until use.

### Cultivation conditions and media

Both strains were inoculated in YP-M medium and in brewing wort containing 138 ppm free amino nitrogen, 0.11 (%w/w) fructose, 4.85 (%w/w) glucose, 11 (%w/w) maltose and 3 (%w/w) maltotriose and allowed to ferment for five days at 16°C. Samples were analyzed after the first and second day of fermentation as these days showed the most important phenotypic manifestation in form of carbon dioxide production. It is important to note that in this work we use gene nomenclature *AGT*1/*MAL*11 and *MTT*1/*MTY*1 unambiguously.

### Inspection and pair-based comparisons of permeases present in the genomic and transcriptomic data of the studied yeasts

We used previously generated genomics (27) and transcriptomic data (28) to collect information about the maltose and maltotriose transporters in strains Sp820 and Sp790. We searched for the transporter genes in each strain in the genome assemblies and annotation files. In this phase, we used the bedtools version 2.30.0 (66) software, UNIX utilities and R custom script to get the gene plot and *loci* information. For resulted permeases, we applied multiple sequence alignment and got the identity matrix for pair-based analysis with *plot_matrix_heapmap.hs* present in GET- HOMOLOGUES-EST (67).

### Permease structure analysis

The structural analysis was done by using the procedure to obtain the amino acids in the maltose translocation mechanism reported earlier (33). Multiple alignment-based sequence conservation analysis was performed using the msa package version 1.32.0 to identify such amino acids in the *MAL* genes of both yeasts (68). Then we obtained the phylogenetic distribution of each permease using maximum likelihood method present in ape version 5.7-1 (69) and phangorn version 2.11.1 packages (70). The topology of transmembrane alpha-helix domains (TMD) in the *MAL* permeases of the studied strains was determined with CCTOP software (71). The 3D model of the *MAL* permeases were built using the AlphaFold DB methodology (72, 73).

### *Mal* genes regulatory region analysis

We used YEASTRACT to identify putative binding sites of the transcription factors MAL63p and Mig1p for each *Mal* upstream regulatory region. (74). According to a previous scientific model (26, 27), at the absence of binding sites to the activator of *Mal* genes, *MAL*63, shows reduced transcription levels for reporter genes or low growth in culture media with maltotriose as the sole carbon source and vice versa.

### FPKM values of the *Mal* genes found in each strain

We proceeded from the previously generated transcriptomic data (28) to obtain the FPKM (Fragment per Kilobase of transcript per Million of mapped reads) values of the transporters *Mal*x1, *AGT*1, *MTT*1, and *MPH*x present in the transcriptome of studied strains, aimed to elucidate the MAL gene expression. Briefly, we generated transcriptomic data from the second and fifth day of fermentation and analyzed it as previously reported (28). Because of either the hybrid nature of the studied yeast, possible specific recombination events, the lack of reliable reference genome and to reproduce previous results (28), we performed a *de novo* assembly.

The transcriptomic pair end reads of both yeasts strain was quality trimmed using Trimmomatic version 0.38 (parameters LEADING:5 TRAILING:5 MINLEN:50) (75) and Sortmerna for contaminant RNA removal (76), then forward and reverse reads for each strain were concatenated independently and assembled using Trinity tool version 2.14.0 with default parameters (77, 78). We annotated the assembled transcripts using Trinotate tool version 3.1.0 (79). For mapping trimmed reads to *de novo* assembled transcript of each strain, Bowtie2 tool version 2.4.4 with default parameters (80) was used. RSEM software version 1.3.3 with default values was used to determine the expression values, as FPKM values, for transcriptome of each yeast strain. From here, the term “isoform” will be implemented in the context of trinity assembly output and left the biological significance about alternative splicing.

### Global expression analysis

Differential gene expression analysis of both yeast strains was carried out using data from samples collected on the second and fifth days of fermentations. To identify statistical differences in gene expression affecting *Mal* genes, we performed a new analysis by read alignment with the reference genome of the yeast *Saccharomyces pastorianus* CBS 1483, (genome and annotations available at NCBI under Bioproject PRJNA522669) using the alignment tool STAR version 2.7.3a with the default parameters (81), to prove that if there are statistical differences in gene expression affecting *MAL* genes. HTSeq-count version 0.11.1 (82) tool was used to convert the BAM alignment files to read count matrices and differential expression analysis was then carried out in edgeR version 3.34.0 (83, 84) filtering genes with low expression level (CPM>5) and contrasting the groups of the second (F2) day of fermentation of both strains (contrast group F2_820-F2_790) and in the same way with the fifth (F5) day of the fermentation (contrast group F5_820-F5_790). A gene significant expression was considered with a threshold of at least twice its abundance in each strain (log2(2) = 1) with a FDR of less than 0.05. From the list of differentially expressed genes, we focus on the *MAL* genes.

### Cellular transport rate test

To determine the transport rate of pNP-α-glucose (structurally related to maltose) and pNP-α-Maltose (structurally related to maltotriose) in our studied strains we performed a cellular transport experiment in standard conditions as previously described (44). The experiments were done in triplicate and controls of previously boiled cells were used. The statistical test, t- Student, was carried-out in R version 4.2.3.

### DNA and RNA extraction

DNA purification was carried out as previously mentioned (86). For RNA extraction, the hot phenol protocol was performed (87), with the only difference that the AES solution (50 mM sodium acetate, 10 mM EDTA, and 0.5% SDS) was used instead of the proposed TES solution.

### Molecular detection of permeases in each strain

For the detection of gene encoding to *Mal*x1, *AGT*1, *MTT*1, and *MPH*3 permeases in both genomes, primers and PCR experiments were performed as described previously (24). A commercial reverse transcriptase (M-MLV Reverse Transcriptase, Promega) was used to detect *MAL* transcripts, following the manufacturer instructions, and using the same PCR conditions as mentioned earlier (24).

### Wort and alpha glycoside consumption analytics

Wort samples were analyzed by HPLC Series 1200 Agilent Technologies using Infrared detector (IR). The mobile phase was H2SO4 (5mM). The flow rate was established to 1 mL per minute using carbohydrates column Aminex HPX-87C Hi-Plex Na, 300 x 7.7 mm and the column was equilibrated as described previously (24). The experiments were done in triplicated and t-Student test was done for statistical significance in the difference of maltose and maltotriose uptake means.

## ACKNOWLEDGMENTS

We thank Viktor Boer (HEINEKEN Research & Development) and Toni Gabaldón (Institute for Research in Biomedicine, Barcelona) for critical inputs and reading of the manuscript. Also, we thank to “Cervecería Cuauhtémoc Moctezuma” for kindly providing us the yeast strains and wort used in this work. E.R.P.O, L.C.D.B, B.P.A and C.I.H.V conceived general idea and project administration. J.H.G.G, and C.I.H.V conceived and designed experiments. C.I.H.V, J.H.G.G and A.G.M.S performed the experiments. B.P.A, J.H.G.G and C.I.H.V analyzed the data. B.P.A, J.H.G.G and C.I.H.V wrote the paper.

## DATA AVAILABILITY STATEMENT

Data that support the findings reported in this work will be made available upon request.

## FOUNDING

## CONFLICT OF INTEREST

The authors declare no conflict of interest.

## SUPPLEMENTARY FIGURES AND TABLES

**Fig. S1.**
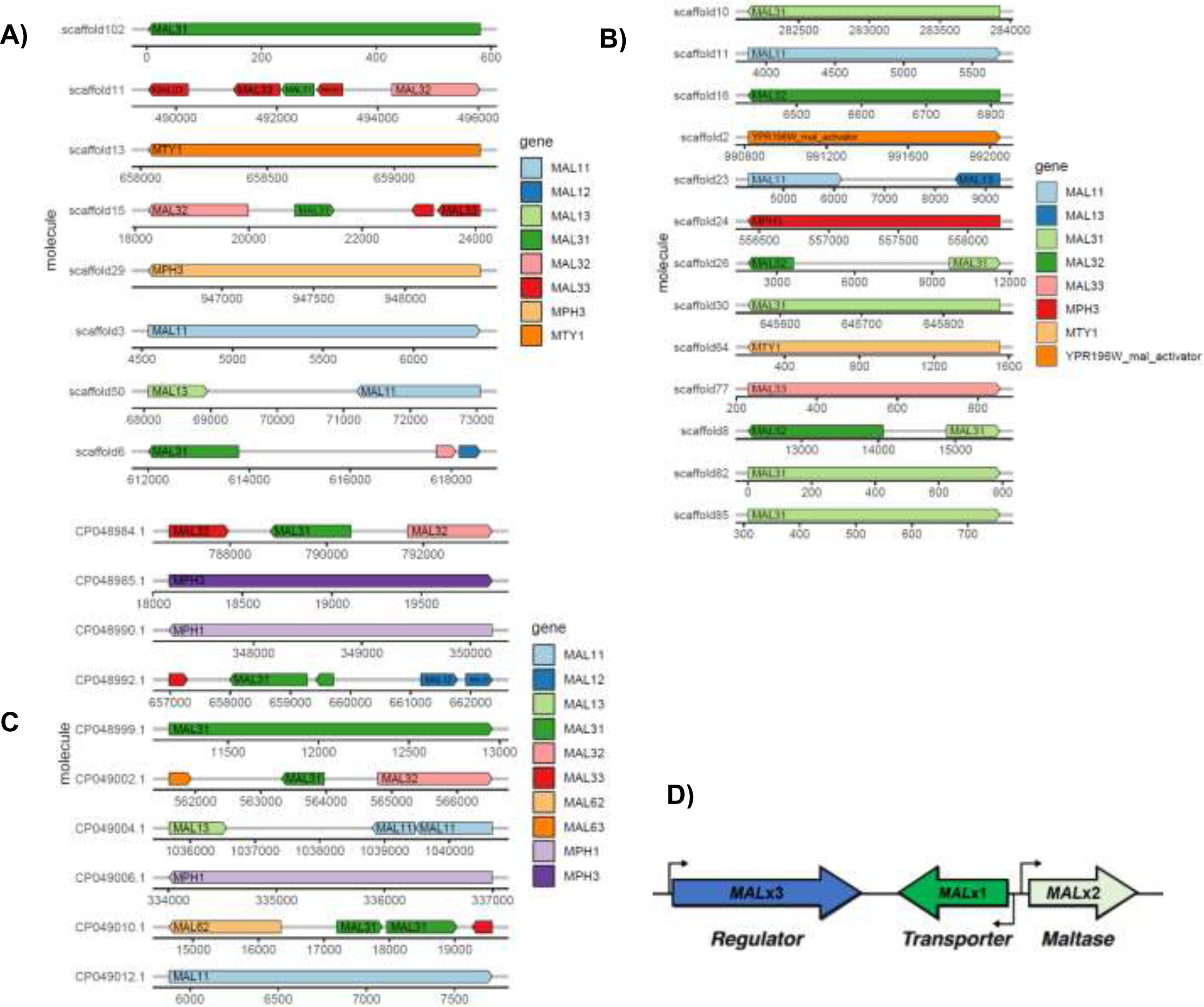
Loci of MAL genes detected in **A)** Sp790 and **B)** Sp820. Notice that not canonic MAL *loci* structure was observed. **C)** MAL *loci* for *Saccharomyces pastorianus* 1483 (NCBI assembly ID: GCA_011022315.1). **D)** Canonical MAL loci modified from earlier report (1). Letter x denotes the locus number where x = 1,2,3,4,6.

**Fig. S2.**
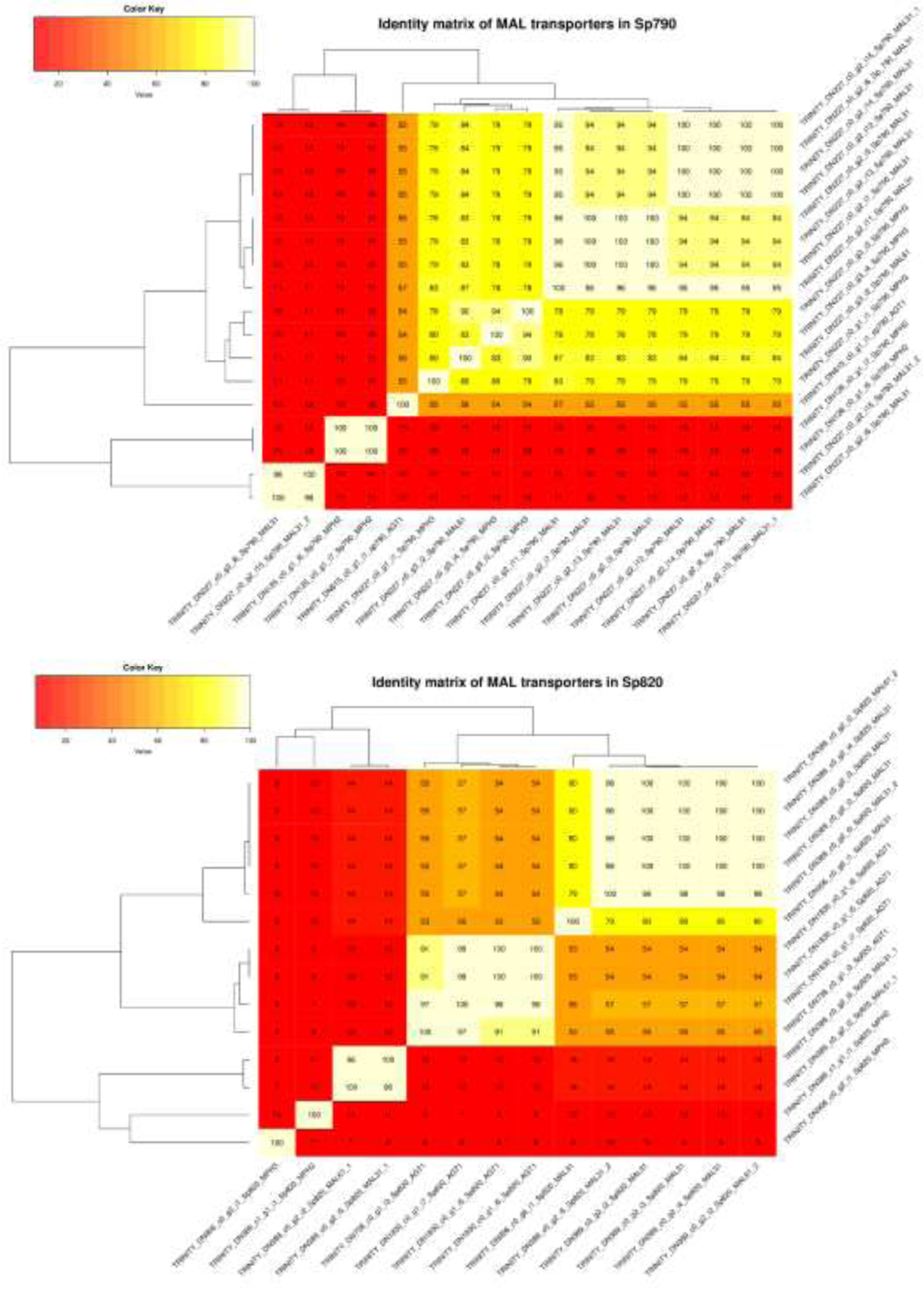
Identity matrix in transporter comparison within each strain. Transcript isoform showing 98% identity with another isoform was taking as same transcript.

**Fig. S3.**
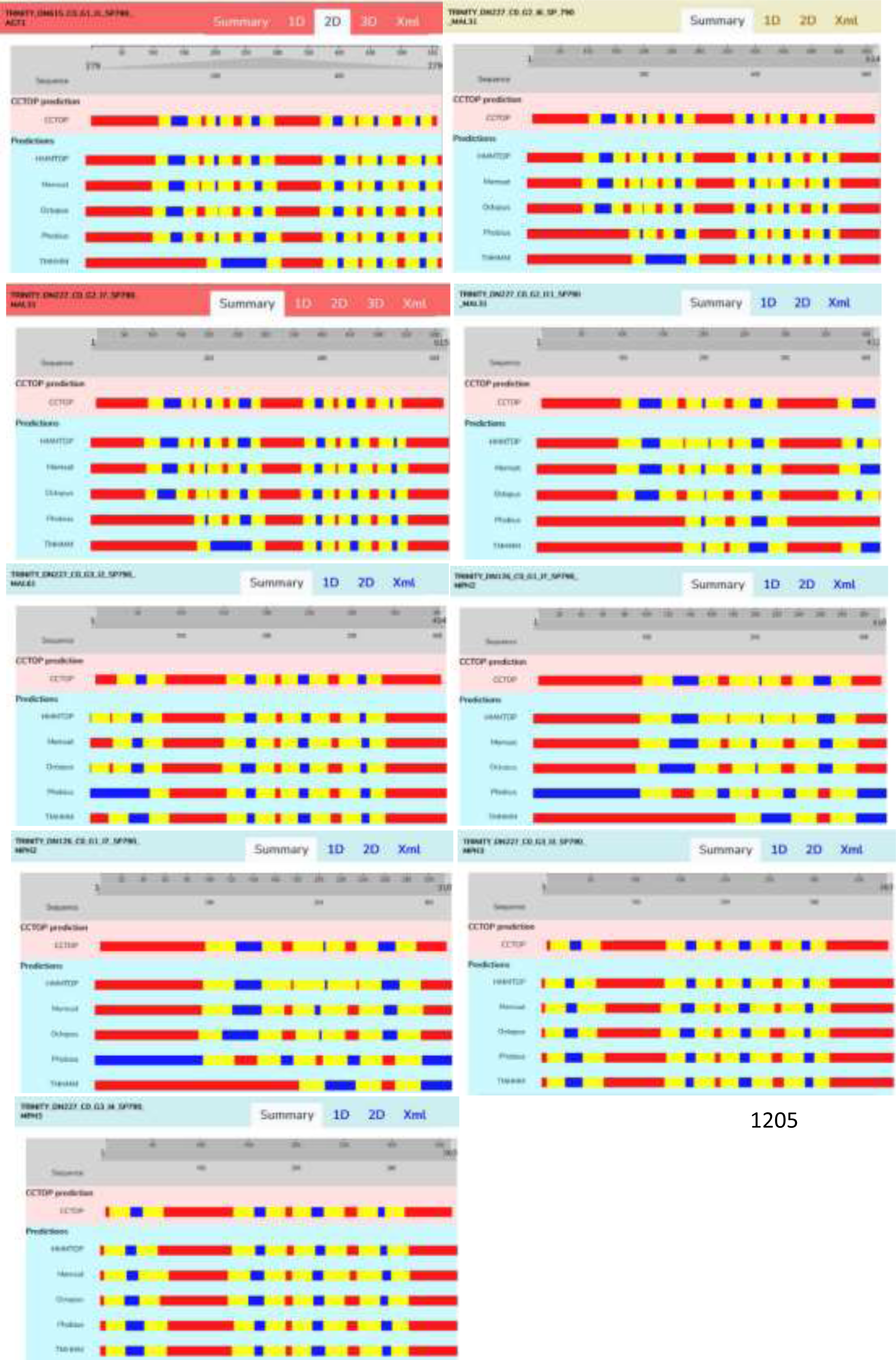
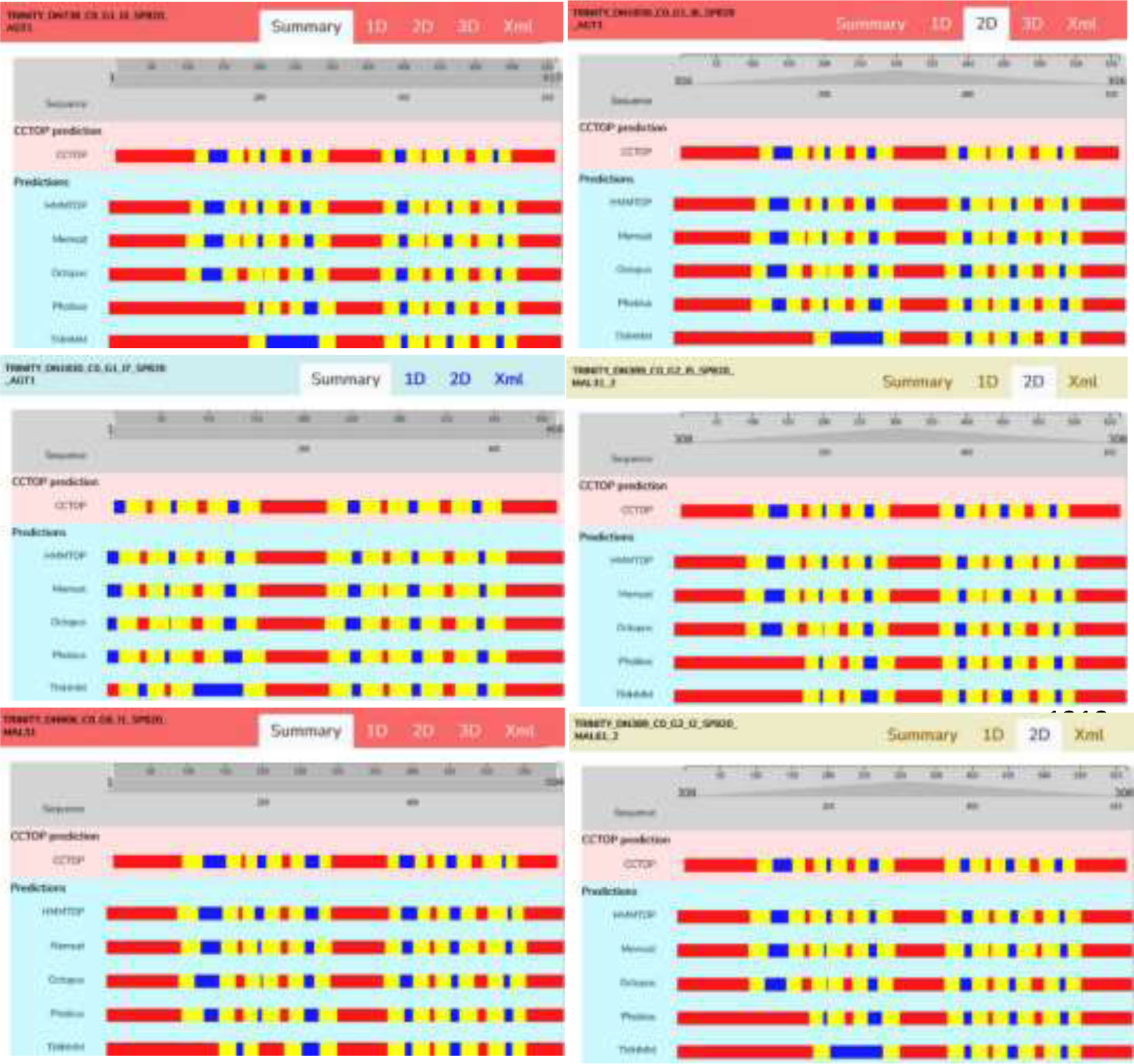
CCTOP transmembrane domain (TMD) models for transcript isoforms annotated as maltose or maltotriose transporter found in each strain. Color-coded amino acid indicate: red: inside/cytosolic; yellow: membrane; blue: outside/extra-cytosolic region. Isoforms TRINITY_DN227_c0_g2_i15_Sp790_MAL31_2 and TRINITY_DN227_c0_g2_i6_Sp790_MAL31 and isoforms TRINITY_DN389_c0_g2_i5_Sp820_MAL31_1, TRINITY_DN389_c0_g2_i2_Sp820_MAL61_1, TRINITY_DN389_c1_g1_i1_Sp820_MPH2 and TRINITY_DN906_c0_g2_i1_Sp820_MPH3 shown no CCTOP transmembrane domain predictions.

**Fig. S4.**
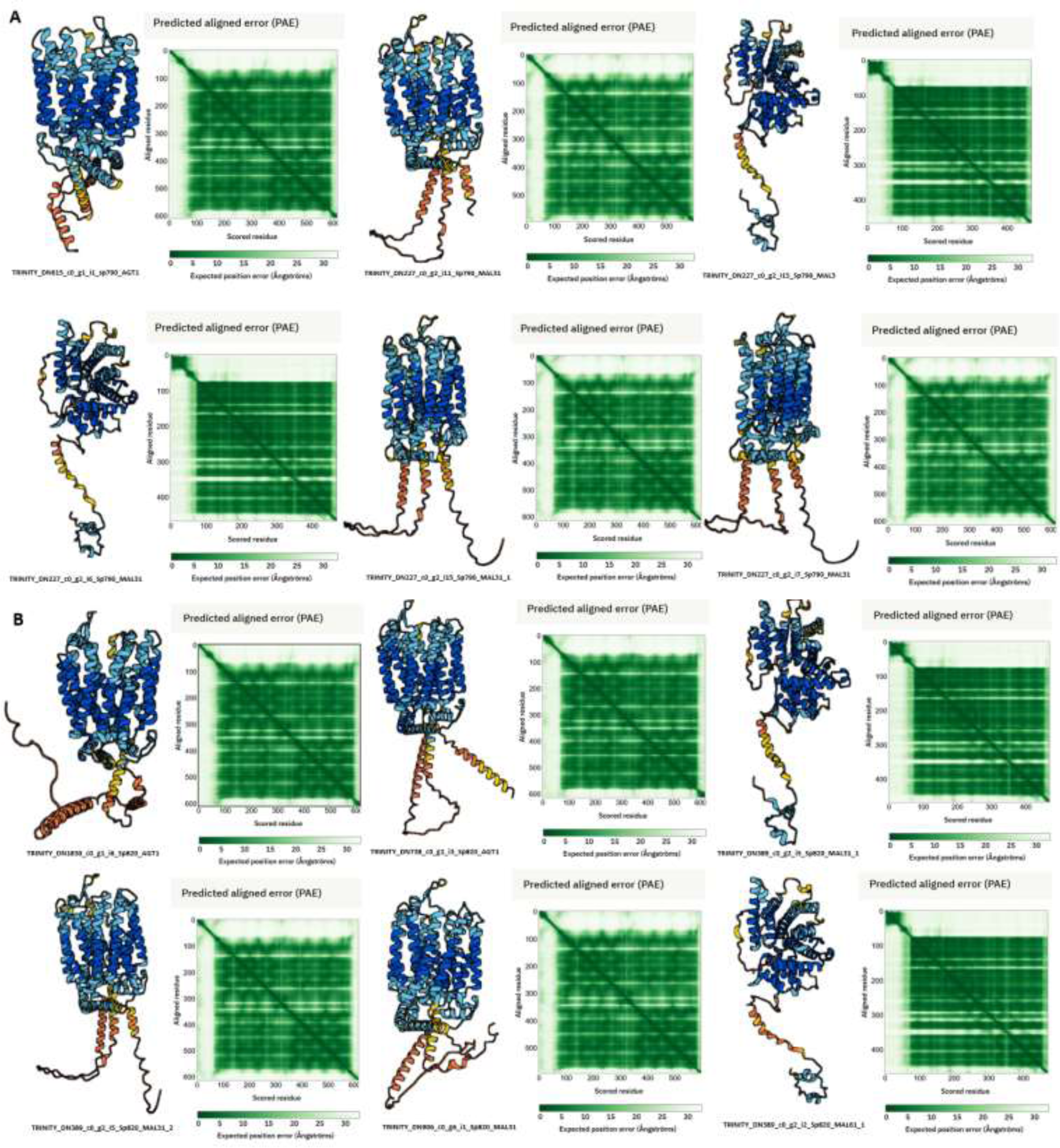
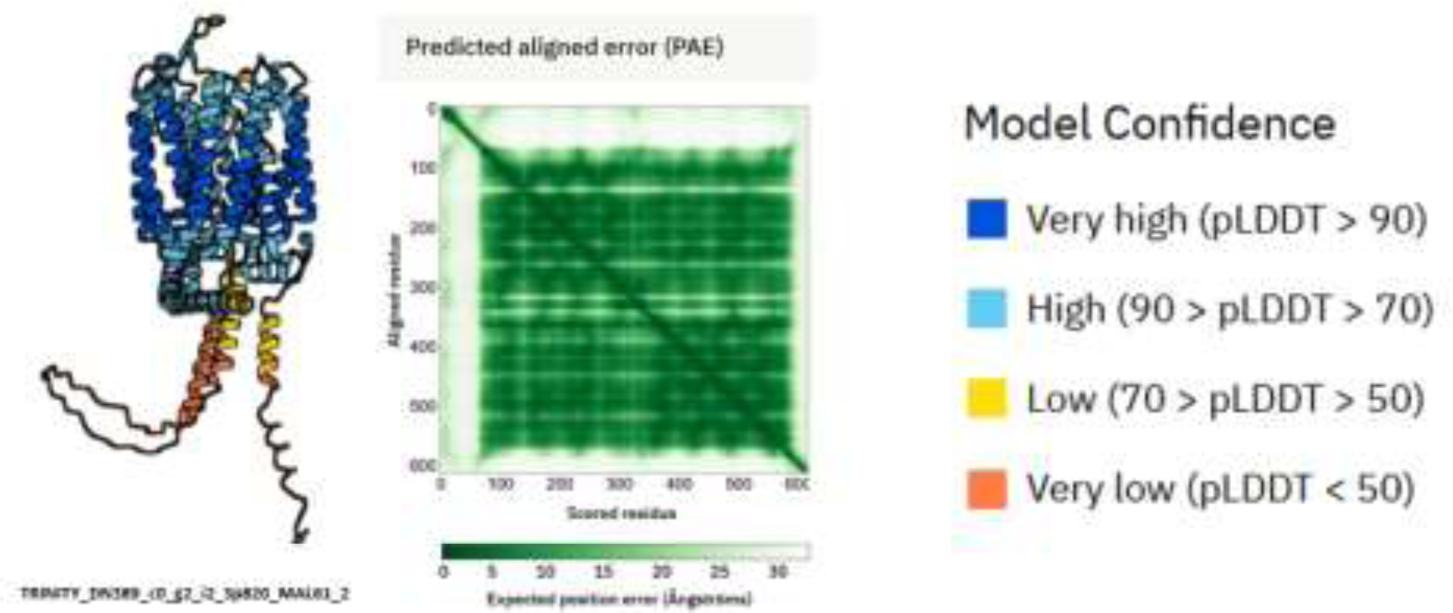
3D structure modeling for non-redundant permeases in for each strain. **A)** 3D model for the permeases presents in Sp790 transcriptome and **B)** 3D model for the permeases present in Sp820 transcriptome. Predicted aligned error plots show the confidence of relative position, in terms of distance depicted in Ångströms, of two residues within the structure predictions. Model confidence, valid for all 3D predictions, indicate local accuracy deduced from its score of Local Distance Difference Test (2, 3).

**Fig. S5.**
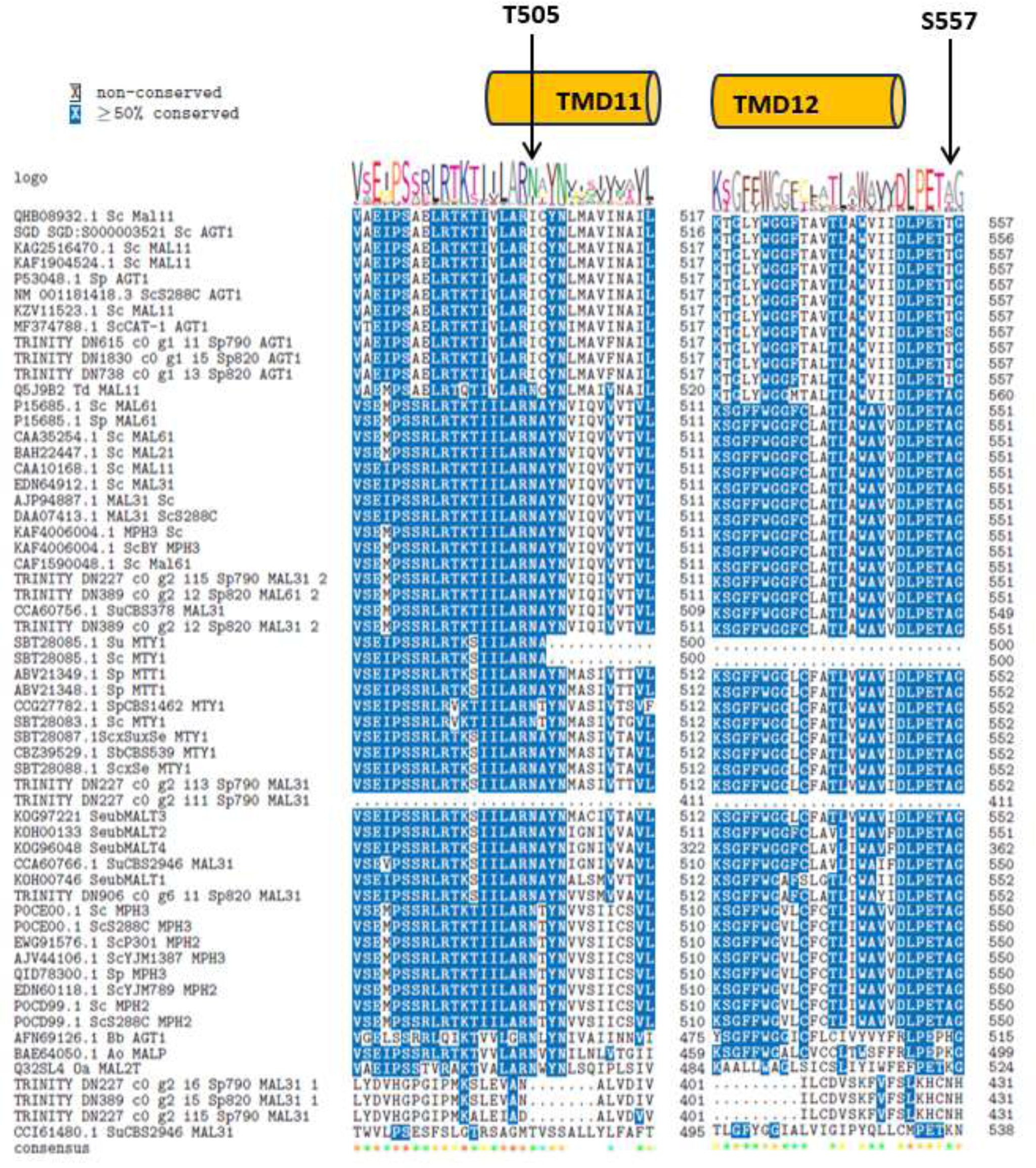
Conservation analysis according to principal insights of Smit and cols (2008) (4). Notice that amino acids T505 and S557 are not conserved throughout the sequences included in this study. Instead of T505 we observed an Arginine (N) conserved above 50% in all transmembrane domain 11 (TMD11) sequences. In the cases of MAL11_i1, from Sp790 and MAL11_i6/i3 we observed an Isoleucine (I) instead of N505 and T505. On the other hand, we observed an Alanine (A) conserved above 50% in all TMD12 sequences. However, for the same permeases mentioned above, we observed a T557 substitution instead of S557 in TMD12.

**Fig. S6.**
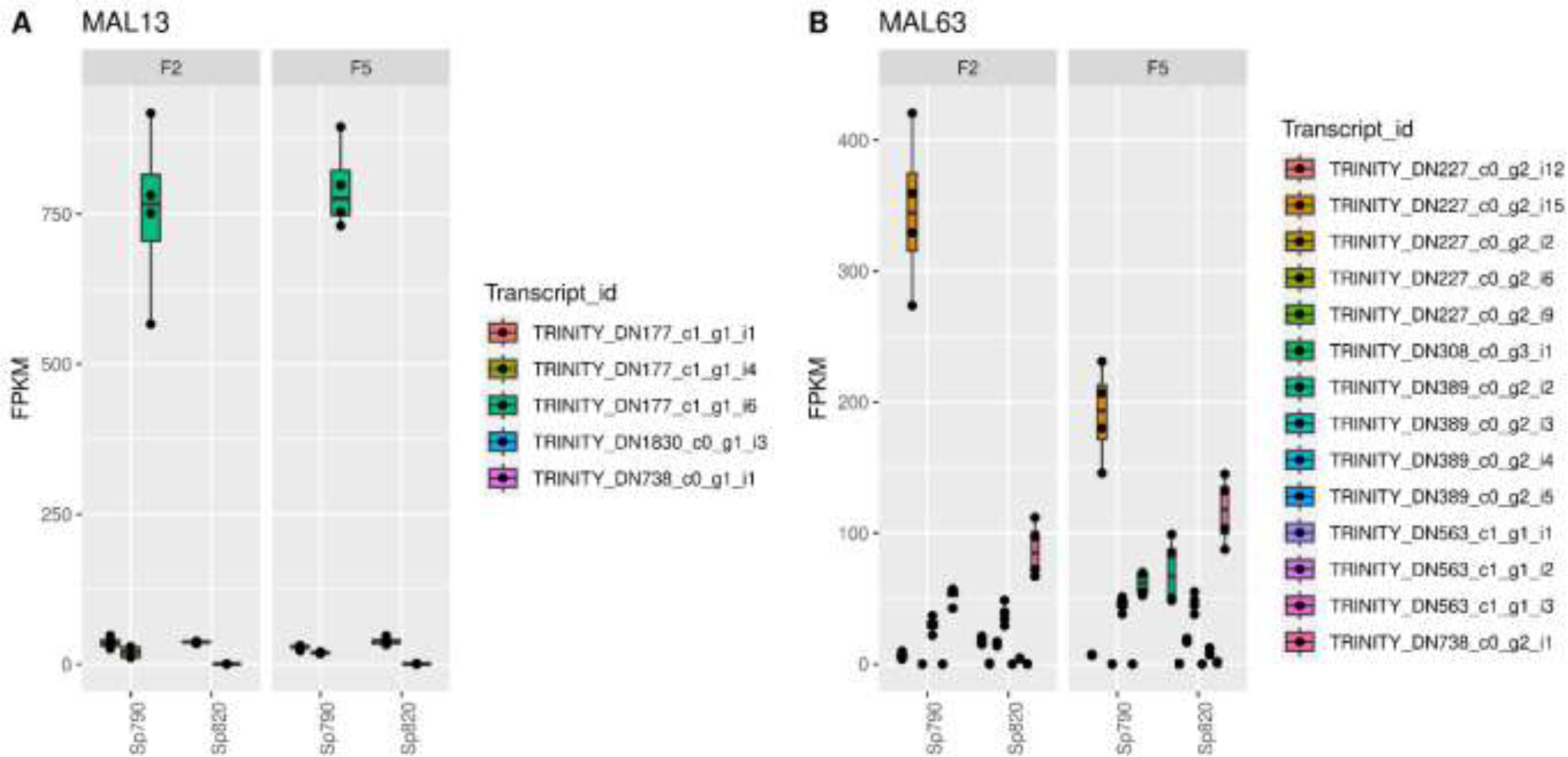
Expression level, in FPKM, of regulatory Mal genes in each strain. **A)** FPKM values for transcript isoforms annotated as *MAL*13 and **B)** for transcripts isoforms annotated as *MAL*63. Note that in both genes Sp790 shows major expression values. F2 and F5 represent the second and fifth day of fermentation, respectively.

**Fig. S7.**
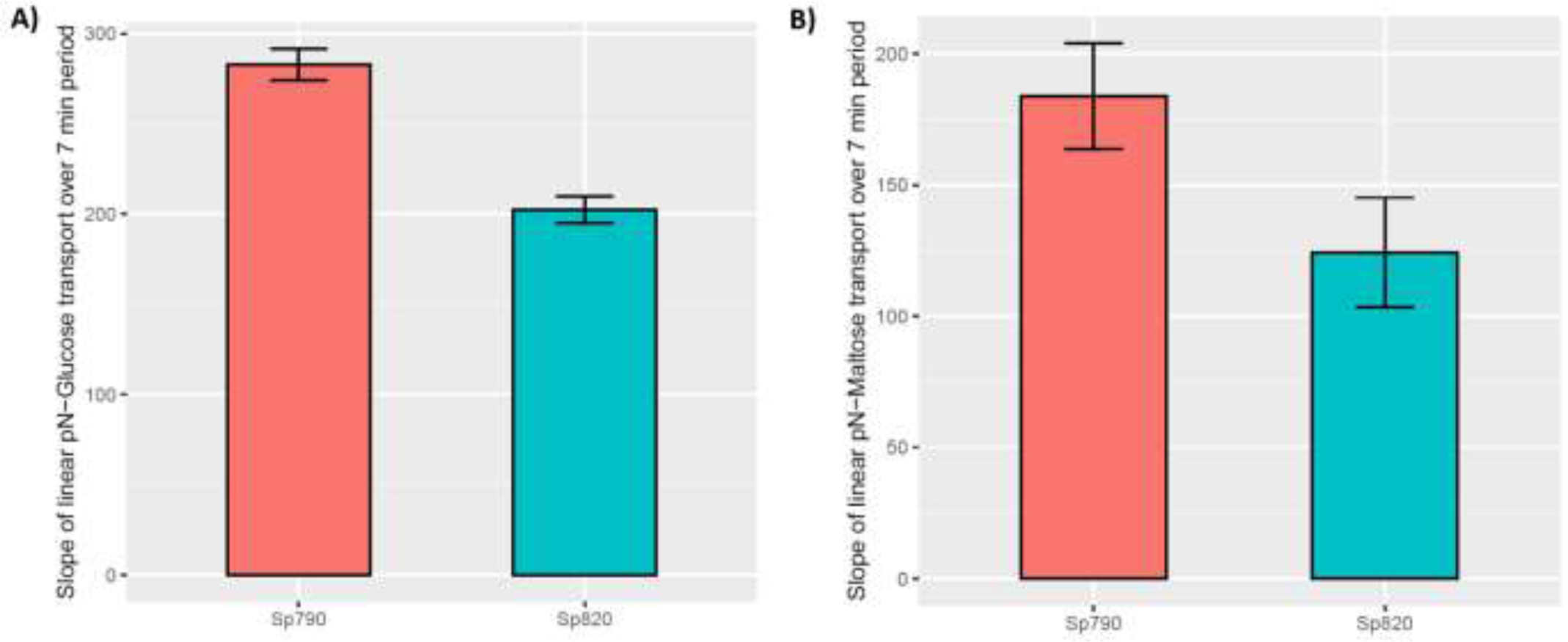
The transport rate of **A)** *para*-nitrophenyl-*α*-d-glucose and **B)** *para*- nitrophenyl-*α*-d-maltose (pN-Glucose and pN-Maltose) uptake by Sp820 and Sp790 strains. The transport rate is determined from the slope of the linear uptake of pNP(G/M) over a 7-min period and normalized to one gram of dry weight. The Strain Sp790 has more transport rate of both pN-glucose and pN-maltose (p = 0.007558 and p = 0.02506, respectively).

**Fig. S8.**
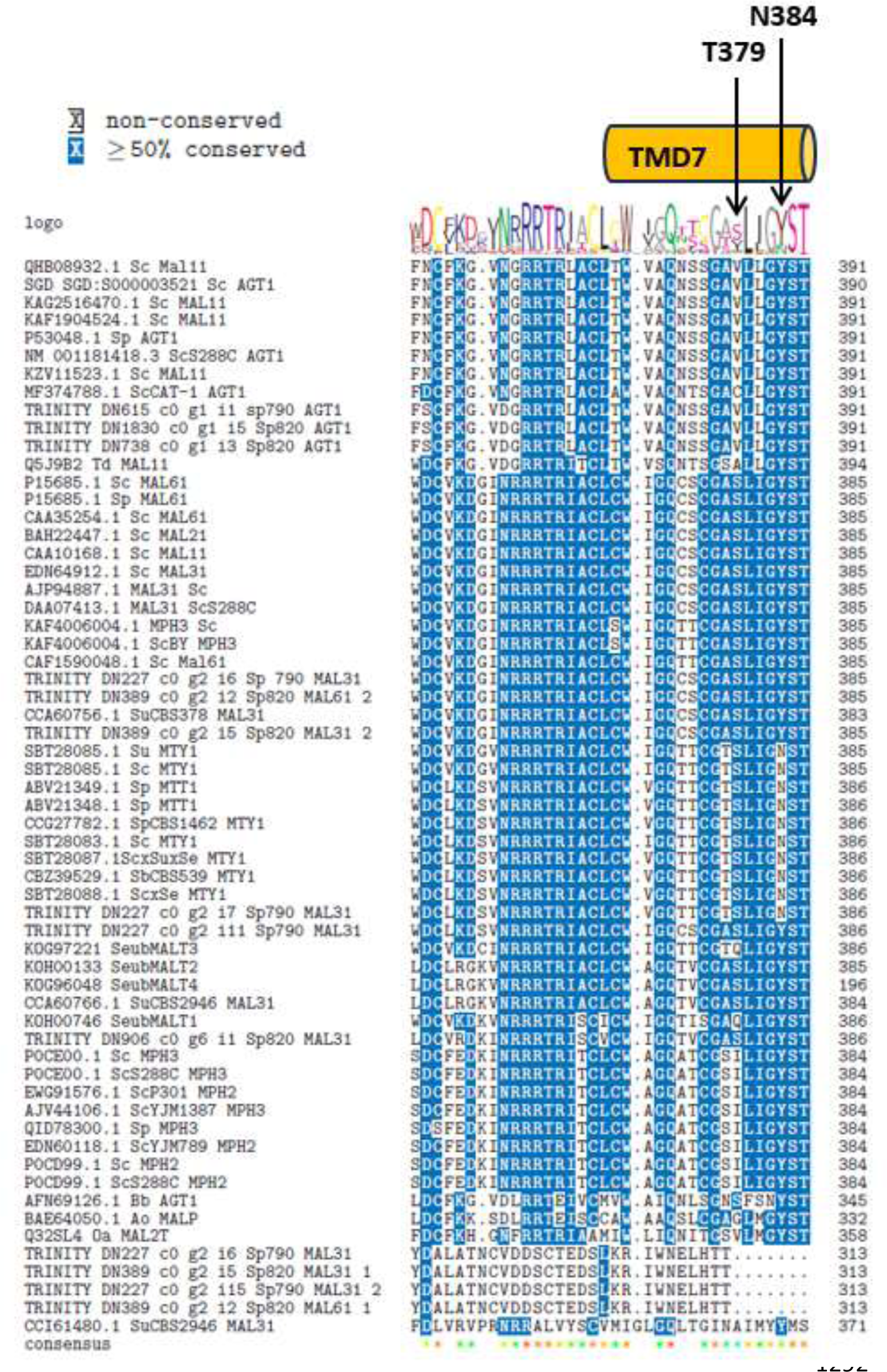
Amin acid conservation analysis for such residues reported as very important for maltotriose substrate specificity in maltose and maltotriosa transporters (5). Note that transcript TRINITY_DN227_c0_g2_i7 from Sp790 has the T379 and N384 conserved as the *MTT*1/*MTY*1 transporters from database.

**Table S1.**
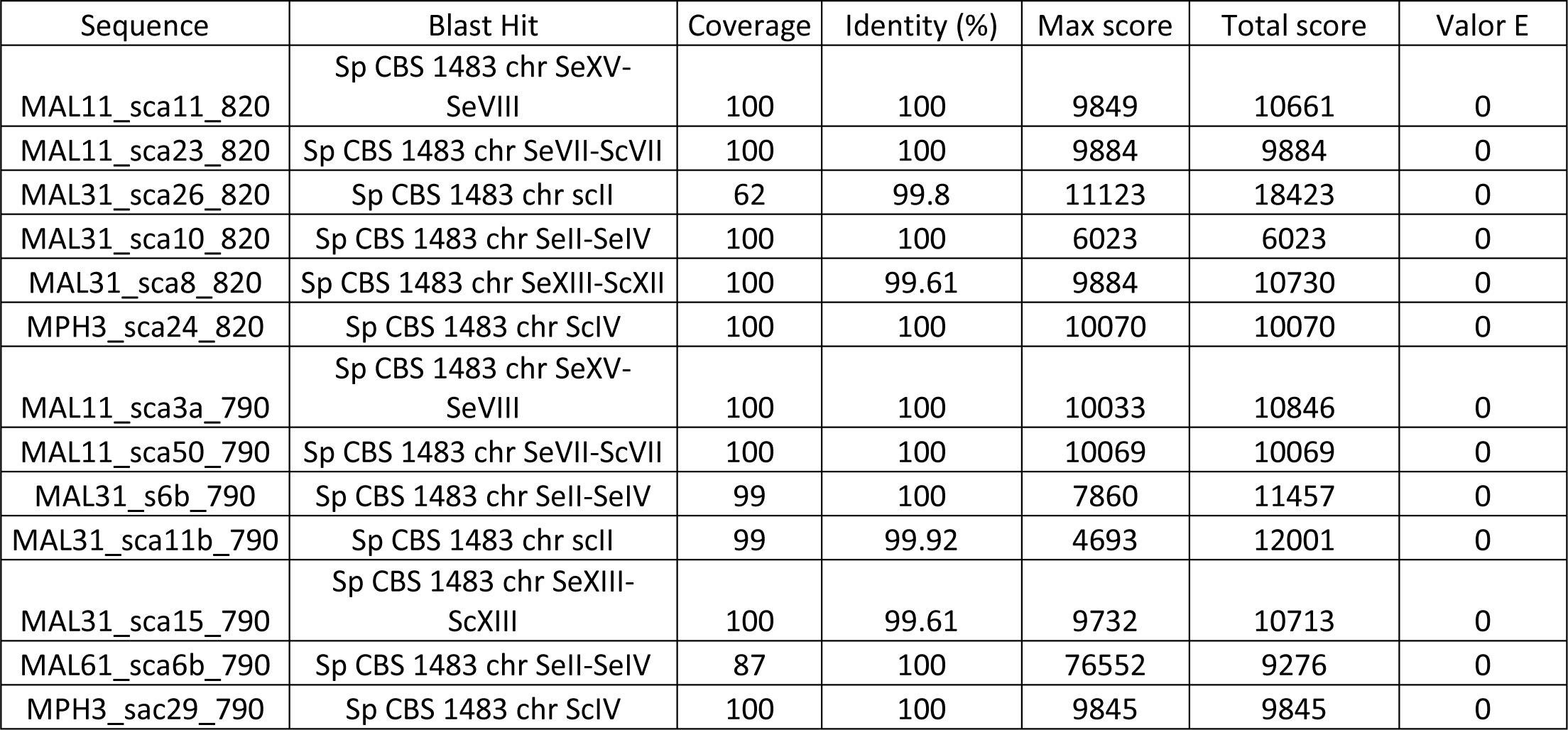
Percentage identity between *Mal loci* of Sp790 and Sp820 with Sp CB1483 *Mal loci*.

**Table S2.** MAL63p and MIG1p binding sites for Sp820.

| Sequence | Gene (aa) | Strand | MAL63 site | Distace from ATG |
| --- | --- | --- | --- | --- |
| MAL11_sca11_reg_820 | AGT1 (610) | "+" | MGCN{9}MGS | -3094 |
|  |  |  | MGCN{9}MGS | -1712 |
|  |  |  | MGCN{9}MGS | -809 |
|  |  |  | MGCN{9}MGS | -799 |
|  |  |  | MGCN{9}MGS | -375 |
|  |  |  | CGGN{9}CGG | -97 |
| MAL11_sca23_reg_820 | AGT1 (394) | "+" | MGCN{9}MGS | -2653 |
|  |  |  | MGCN{9}MGS | -2623 |
|  |  |  | MGCN{9}MGS | -1001 |
| MAL31_sca8_reg_820 | MAL31 (231-383) | "+" | MGCN{9}MGS | -538 |
|  |  |  | CGGN{9}CGC | -646 |
| MAL31_sca10_reg_820 | MAL31 (594) | "-" | NA | NA |
| MPH3_sca24_reg_820 | MPH3 (602) | "-" | CGGN{9}CGG | -551 |
| Sequence | Gene (aa) | Strand | Mig1p sites | Distance from ATG |
| MAL11_sca11_reg_820 | AGT1 (610) | "+" | CCCCRC | -573 |
|  |  |  | CCCCRC | -564 |
|  |  |  | SYGGRG | -1054 |
| MAL11_sca23_reg_820 | AGT1 (394) | "+" | CCCCRC | -1119 |
| MAL31_sca8_reg_820 | MAL31 (231-383) | "+" | CCCCRC | -735 |
|  |  |  | SYGGRG | -1018 |
|  |  |  | SYGGRG | -781 |
| MAL31_sca10_reg_820 | MAL31 (594) | "-" | CCCCRC | -432 |
|  |  |  | SYGGRG | -456 |
|  |  |  | SYGGRG | -472 |
|  |  |  | SYGGRG | -761 |
| MPH3_sca24_reg_820 | MPH3 (602) | "-" | CCCCRC | -2809 |
|  |  |  | SYGGRG | -923 |
|  |  |  | SYGGRG | -2741 |
|  |  |  | SYGGRG | -2928 |

**Table S3.** MAL63p and MIGip binding sites for Sp790.

| Sequence | Gene (aa) | Strand | MAL63 site | Distance from ATG |
| --- | --- | --- | --- | --- |
| MAL11_sca3a_reg_790 | AGT1 (610) | "+" | MGCN{9}MGS | -3014 |
|  |  |  | MGCN{9}MGS | -1632 |
|  |  |  | MGCN{9}MGS | -729 |
|  |  |  | MGCN{9}MGS | -719 |
|  |  |  | MGCN{9}MGS | -295 |
|  |  |  | CGGN{9}CGG | -17 |
| MAL11_sca50_reg_790 | AGT1 (394) | "-" | MGCN{9}MGS | -67 |
|  |  |  | MGCN{9}MGS | -604 |
|  |  |  | MGCN{9}MGS | -634 |
|  |  |  | MGCN{9}MGS | -2256 |
| MAL31_sca6b_reg_790 | MAL31 (594) | "-" | NA | NA |
| MAL31_sca11b_reg_790 | MAL31 (168-261-217) | "-" | MGCN{9}MGS | -571 |
|  |  |  | MGCN{9}MGS | -669 |
|  |  |  | MGCN{9}MGS | -796 |
|  |  |  | MGCN{9}MGS | -797 |
|  |  |  | MGCN{9}MGS | -798 |
|  |  |  | MGCN{9}MGS | -799 |
|  |  |  | MGCN{9}MGS | -800 |
|  |  |  | MGCN{9}MGS | -801 |
|  |  |  | MGCN{9}MGS | -802 |
|  |  |  | MGCN{9}MGS | -803 |
|  |  |  | MGCN{9}MGS | -804 |
|  |  |  | MGCN{9}MGS | -805 |
|  |  |  | MGCN{9}MGS | -806 |
|  |  |  | MGCN{9}MGS | -807 |
|  |  |  | MGCN{9}MGS | -808 |
|  |  |  | MGCN{9}MGS | -813 |
|  |  |  | MGCN{9}MGS | -818 |
|  |  |  | MGCN{9}MGS | -1464 |
|  |  |  | MGCN{9}MGS | -1686 |
|  |  |  | MGCN{9}MGS | -1688 |
|  |  |  | MGCN{9}MGS | -1689 |
|  |  |  | MGCN{9}MGS | -1690 |
|  |  |  | MGCN{9}MGS | -1691 |
|  |  |  | MGCN{9}MGS | -1692 |
|  |  |  | MGCN{9}MGS | -1693 |
|  |  |  | MGCN{9}MGS | -1694 |
|  |  |  | MGCN{9}MGS | -3110 |
| MAL31_sca15_reg_790 | MAL31<br>(231-383) | "+" | MGCN{9}MGS | -596 |
|  |  |  | CGGN{9}CGC | -704 |
| MPH3_sca | MPH3<br>(602) | "-" | CGGN{9}CGG | -308 |
| Sequence | Gene (aa) | Strand | Mig1p site | Distance from ATG |
| MAL11_sca3a_reg_790 | AGT1 (610) | "+" | CCCCRC | -493 |
|  |  |  | CCCCRC | -484 |
|  |  |  | SYGGRG | -974 |
| MAL11_sca50_reg_790 | AGT1 (394) | "-" | CCCCRC | -2129 |
| MAL31_sca6b_reg_790 | MAL31<br>(594) | "-" | CCCCRC | -719 |
|  |  |  | SYGGRG | -243 |
|  |  |  | SYGGRG | -743 |
|  |  |  | SYGGRG | -759 |
|  |  |  | SYGGRG | -1048 |
| MAL31_sca11b_reg_790 | MAL31<br>(168-261-217) | "-" | SYGGRG | -788 |
|  |  |  | SYGGRG | -789 |
|  |  |  | SYGGRG | -790 |
|  |  |  | SYGGRG | -791 |
|  |  |  | SYGGRG | -792 |
|  |  |  | SYGGRG | -793 |
|  |  |  | SYGGRG | -794 |
|  |  |  | SYGGRG | -795 |
|  |  |  | SYGGRG | -796 |
|  |  |  | SYGGRG | -797 |
|  |  |  | SYGGRG | -798 |
|  |  |  | SYGGRG | -799 |
|  |  |  | SYGGRG | -800 |
|  |  |  | SYGGRG | -801 |
|  |  |  | SYGGRG | -802 |
|  |  |  | SYGGRG | -803 |
|  |  |  | SYGGRG | -804 |
|  |  |  | SYGGRG | -805 |
|  |  |  | SYGGRG | -806 |
|  |  |  | SYGGRG | -807 |
|  |  |  | SYGGRG | -808 |
|  |  |  | SYGGRG | -960 |
|  |  |  | SYGGRG | -1041 |
|  |  |  | SYGGRG | -1678 |
|  |  |  | SYGGRG | -1679 |
|  |  |  | SYGGRG | -1680 |
|  |  |  | SYGGRG | -1681 |
|  |  |  | SYGGRG | -1682 |
|  |  |  | SYGGRG | -1683 |
|  |  |  | SYGGRG | -1684 |
|  |  |  | SYGGRG | -1685 |
|  |  |  | SYGGRG | -1686 |
|  |  |  | SYGGRG | -1687 |
|  |  |  | SYGGRG | -1688 |
|  |  |  | SYGGRG | -1689 |
|  |  |  | SYGGRG | -1690 |
|  |  |  | SYGGRG | -1691 |
|  |  |  | SYGGRG | -1692 |
|  |  |  | SYGGRG | -1693 |
|  |  |  | SYGGRG | -1694 |
|  |  |  | SYGGRG | -3044 |
|  |  |  | SYGGRG | -3112 |
|  |  |  | SYGGRG | -3113 |
|  |  |  | SYGGRG | -3150 |
| MAL31_sca15_reg_790 | MAL31<br>(231-383) | "+" | CCCCRC | -793 |
|  |  |  | SYGGRG | -1076 |
|  |  |  | SYGGRG | -839 |
| MPH3_sca29_reg_790 | MPH3<br>(602) | "-" | CCCCRC | -2566 |
|  |  |  | SYGGRG | -680 |
|  |  |  | SYGGRG | -2498 |
|  |  |  | SYGGRG | -2685 |

**Table S4.**
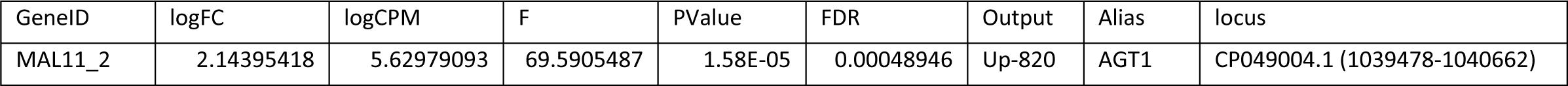
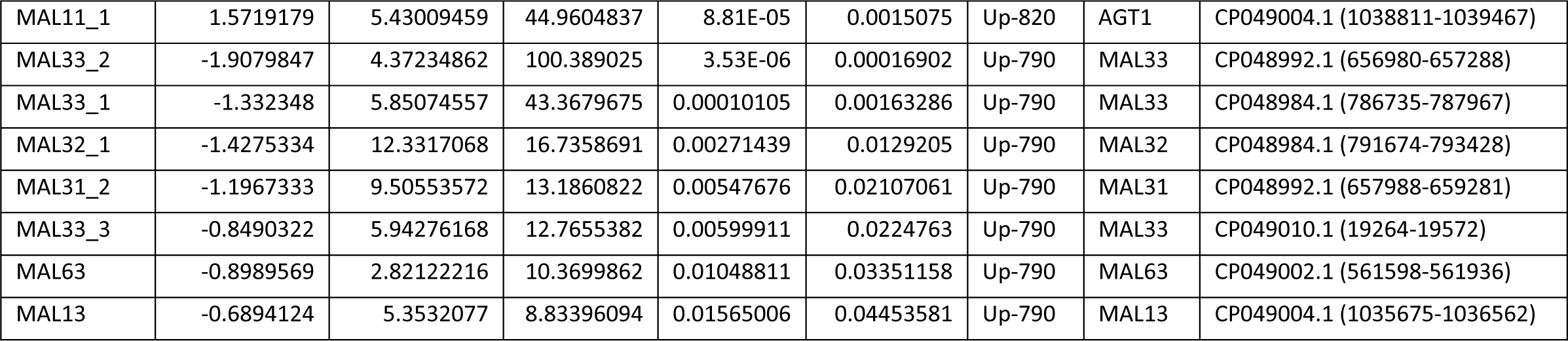
Differential expressed *Mal* genes in Sp790 and Sp820 at the second day of fermentation.

**Table S5.** Differential expressed *Mal* genes in Sp790 and Sp820 at the fifth day of fermentation.

| GeneID | logFC | logCPM | F | PValue | FDR | Output | GeneID | Alias | Locus |
| --- | --- | --- | --- | --- | --- | --- | --- | --- | --- |
| MAL32_2 | 2.19588889 | 6.50332801 | 33.3317798 | 0.00010532 | 0.00084308 | Up-820 | MAL32_2 | MAL32 | CP049002.1 (564788-566537) |
| MAL62 | 1.96025696 | 6.68733696 | 27.4746788 | 0.0002404 | 0.00148702 | Up-820 | MAL62 | MAL62 | CP049010.1 (14632-16342) |
| MAL11_2 | 1.47549449 | 5.72153066 | 26.7079494 | 0.00025771 | 0.00156519 | Up-820 | MAL11_2 | AGT1 | CP049004.1 (1039478-1040662) |
| MAL12_1 | 1.35372616 | 5.66952585 | 20.1485458 | 0.0008277 | 0.0035503 | Up-820 | MAL12_1 | MAL12 | CP048992.1 (661169-661778) |
| MAL12_2 | 1.03114719 | 5.60129636 | 12.4092297 | 0.00447231 | 0.01296753 | Up-820 | MAL12_2 | MAL12 | CP048992.1 (661917-662357) |
| MAL31_6 | 0.70359396 | 3.81192893 | 6.80631706 | 0.02330847 | 0.04825076 | Up-820 | MAL31_6 | MAL31 | CP049010.1 (17197-17892) |
| MAL63 | -2.5881092 | 4.44857299 | 86.5539567 | 9.69E-07 | 3.71E-05 | Up-790 | MAL63 | MAL63 | CP049002.1 (561598-561936) |
| MAL11_3 | -2.3013114 | 5.23431861 | 30.1410846 | 0.00016267 | 0.0011289 | Up-790 | MAL11_3 | AGT1 | CP049012.1 (5881-7713) |
| MAL33_3 | -2.3135934 | 7.82153538 | 29.1297917 | 0.00018805 | 0.00124606 | Up-790 | MAL33_3 | MAL33 | CP049010.1 (19264-19572) |
| MAL33_2 | -1.4274395 | 4.80552978 | 26.9996103 | 0.00024892 | 0.00152276 | Up-790 | MAL33_2 | MAL33 | CP048992.1 (656980-657288) |
| MAL33_1 | -1.1914058 | 6.30857726 | 16.1741444 | 0.00179126 | 0.0063814 | Up-790 | MAL33_1 | MAL33 | CP048984.1 (786735-787967) |
| MAL31_4 | -1.4115258 | 4.93950783 | 15.5937076 | 0.00210243 | 0.00723693 | Up-790 | MAL31_4 | MAL31 | CP048999.1 (11175-12959) |

